# Schwann cell release of p11 induces sensory neuron hyperactivity in Fabry disease

**DOI:** 10.1101/2023.05.26.542493

**Authors:** Tyler B. Waltz, Dongman Chao, Eve K. Prodoehl, Vanessa L. Ehlers, Bhavya S. Dharanikota, Nancy M. Dahms, Elena Isaeva, Quinn H. Hogan, Bin Pan, Cheryl L. Stucky

## Abstract

Patients with Fabry disease suffer from chronic debilitating pain and peripheral sensory neuropathy with minimal treatment options, but the cellular drivers of this pain are unknown. Here, we propose a novel mechanism by which altered signaling between Schwann cells and sensory neurons underlies the peripheral sensory nerve dysfunction we observe in a genetic rat model of Fabry disease. Using *in vivo* and *in vitro* electrophysiological recordings, we demonstrate that Fabry rat sensory neurons exhibit pronounced hyperexcitability. Schwann cells likely contribute to this finding as application of mediators released from cultured Fabry Schwann cells induces spontaneous activity and hyperexcitability in naïve sensory neurons. We examined putative algogenic mediators using proteomic analysis and found that Fabry Schwann cells release elevated levels of the protein p11 (S100-A10) which induces sensory neuron hyperexcitability. Removal of p11 from Fabry Schwann cell media causes hyperpolarization of neuronal resting membrane potential, indicating that p11 contributes to the excessive neuronal excitability caused by Fabry Schwann cells. These findings demonstrate that rats with Fabry disease exhibit sensory neuron hyperexcitability caused in part by Schwann cell release of the protein p11.

**Graphical Abstract:** 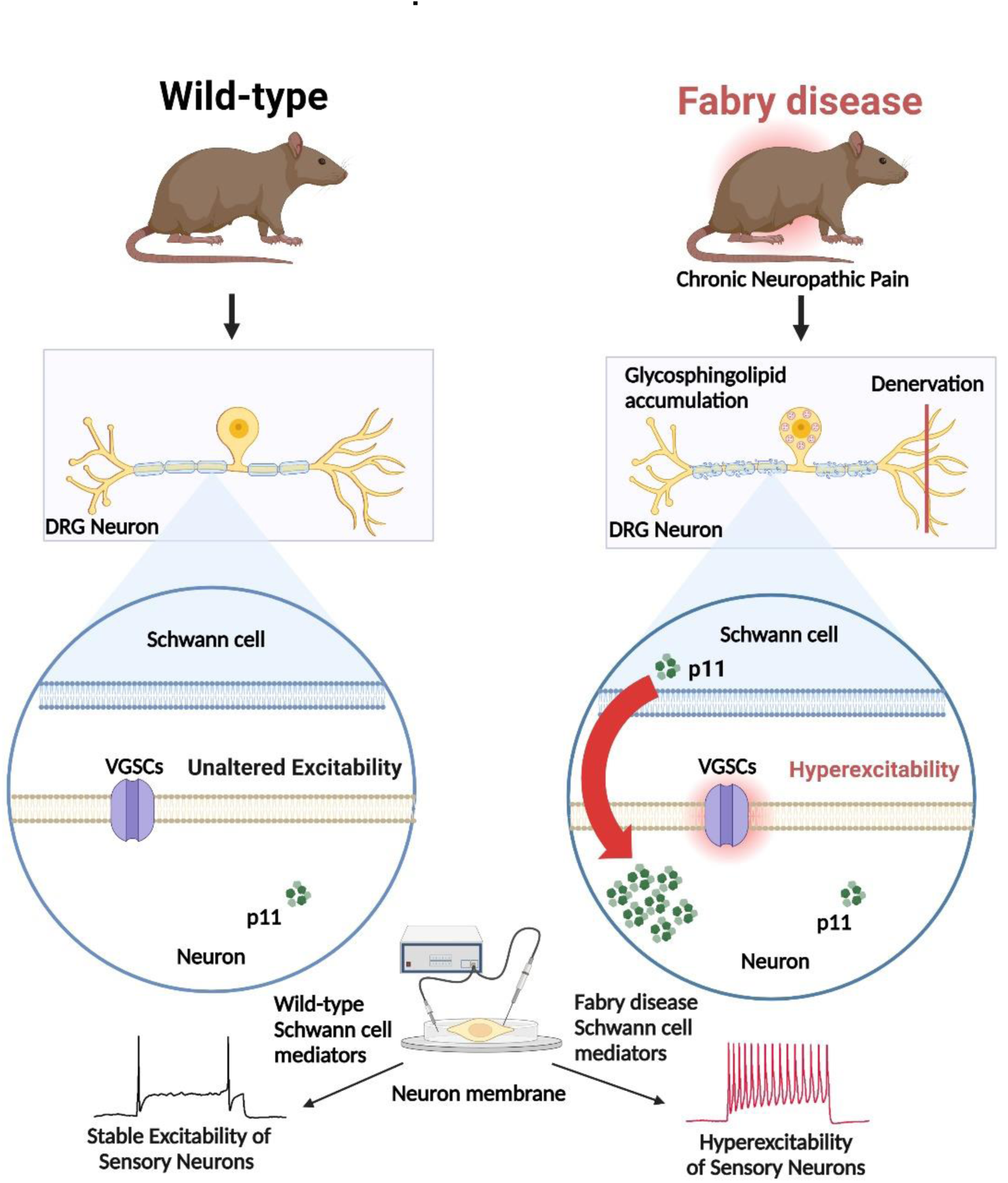

## Introduction

Fabry disease, the most common X-linked lysosomal storage disease, is caused by a deficiency in the enzyme alpha-galactosidase A (α-Gal A), which results in chronic intracellular accumulation of glycosphingolipids in multiple tissues, including sensory dorsal root ganglia (DRG) neurons. Though initially considered a rare genetic disorder, recent studies suggest Fabry disease affects up to 1:1600 individuals [1, 2] and is under-diagnosed [3]. The most debilitating complication is childhood onset of severe chronic pain, described as mechanical allodynia and burning in the hands and feet. Many patients also endure recurring attacks of intense episodic pain that occur spontaneously or are triggered by extreme temperature, fever, fatigue, or stress [4]. Existing treatments for pain are inadequate, and enzyme replacement therapy does not alleviate the pain [5–8]. The pathophysiological mechanisms underlying the chronic and episodic pain in Fabry disease are poorly understood, yet this topic receives little research attention. Fabry pain has neuropathic attributes and patients display evidence of peripheral neuropathy including structural abnormalities in the dorsal root ganglia and peripheral nerves [9–14]. However, it has not been established that pain in Fabry disease is specifically attributable to sensory neuron dysfunction, as pain may be attributed to more central sites, including the spinal cord and brain [15–17]. Furthermore, peripheral nerves from patients with Fabry disease show morphological changes in Schwann cells [11, 13, 18]. Emerging evidence suggests that Schwann cells, which are critical for maintaining healthy nerve function, can induce pain by directly increasing the activity of nociceptive neurons [19–22], although it is still unclear how Schwann cells elicit these effects. Here, we used a recently characterized genetic rat model [23] to investigate how peripheral sensory neurons and Schwann cells may contribute to Fabry disease pain.

## Results

### Peripheral sensory neurons from Fabry rats exhibit spontaneous activity and mechanical sensitization

Rats with completely deficient α-Gal A activity (Fabry rats) [23] exhibit ongoing, spontaneous pain behaviors [24]. We first examined whether this is attributable to spontaneous activity in peripheral sensory neurons, as is commonly the case in other neuropathic pain models as well as human subjects [25–35]. Specifically, we used *in vivo* dorsal root teased fiber electrophysiological recordings in anesthetized rats to determine whether peripheral nerves in Fabry disease exhibit spontaneous activity (Figure 1A). Lumbar 4 (L4) dorsal roots of Fabry and wild-type (WT) rats were teased into fine bundles and the incidence and frequency of spontaneous action potentials were recorded (Figure 1B). Fabry rats displayed a higher proportion of spontaneous afferent activity per bundle (Figure 1C) and per animal (Figure 1D) compared to WT rats. Dorsal root bundles from Fabry rats also exhibited a trend towards increased spontaneous firing frequency (Figure 1E).

**Figure 1:**
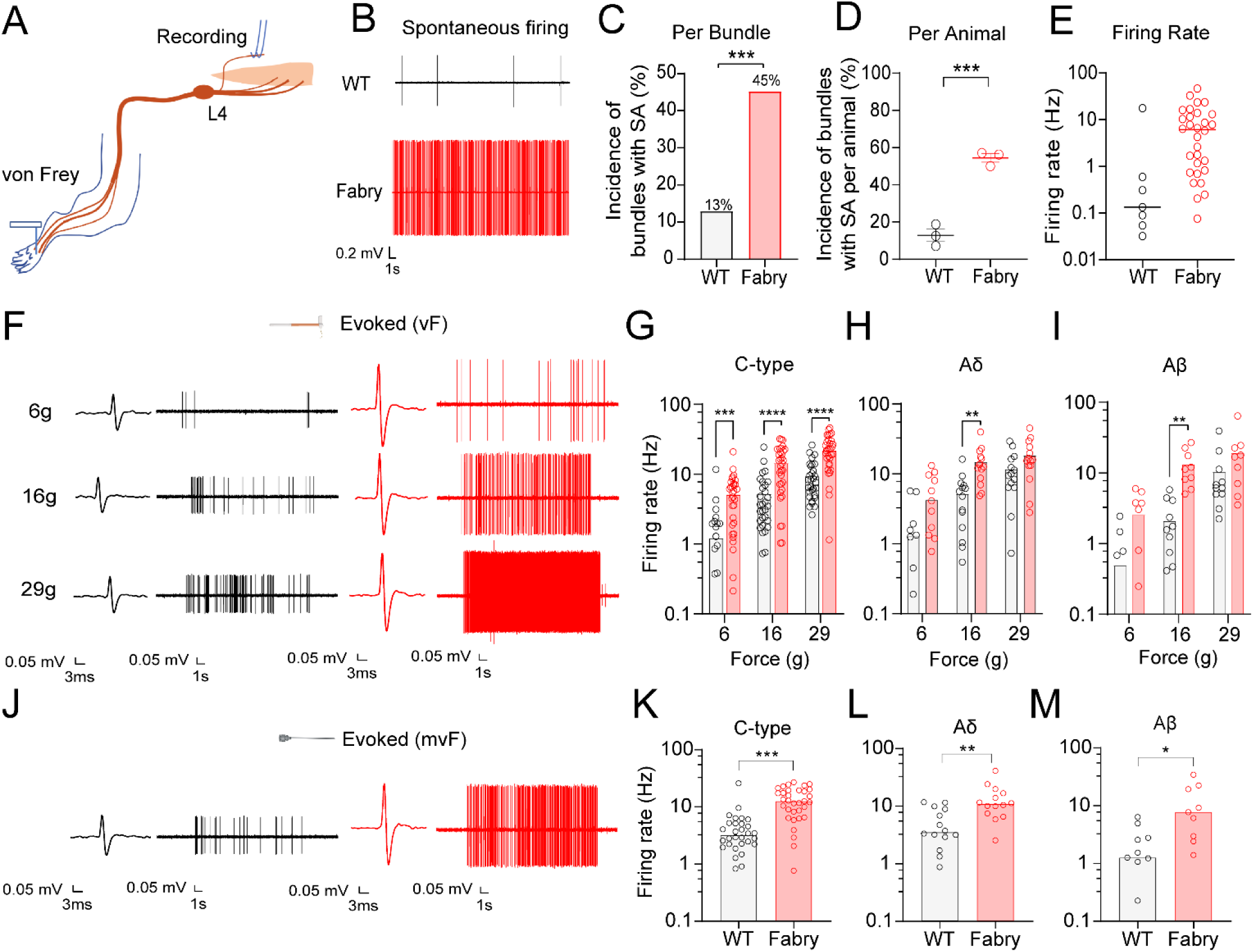
Peripheral dorsal roots from Fabry rats exhibit spontaneous activity and mechanical hypersensitivity. A) L4 dorsal roots from Fabry or WT rats were teased into bundles and recorded without external stimulation or with von Frey stimulation of the hindpaw. B) Representative traces of spontaneously firing dorsal roots from Fabry or WT rats. C) Teased bundles from Fabry rats exhibited a higher proportion of spontaneous activity as analyzed per teased bundle or D) per animal. E) Spontaneous firing frequency between Fabry (red) and WT (black) rats (unpaired *t*-test, p = 0.16). F) Representative traces of recorded teased, single unit C-fiber activity due to innocuous von Frey (vF) stimulation of the plantar hindpaw. Single (G) C-, (H) Aδ, and (I) Aβ units from Fabry rats exhibit increased firing frequency due to graded innocuous von Frey stimulation. J) Representative traces of C-fiber activity due to stimulation of the hindpaw with a modified von Frey filament with a tungsten tip (mvF). (K) C-, (L) Aδ and (M) Aβ units from Fabry rats exhibit increased firing frequency due to noxious mvF stimulation of the hindpaw, Fabry = red, WT = black. n = 3 rats per genotype. Data for (C, G-I) reported as mean; (D) as mean ± SEM; (E, K-M) as median. (C, K-M) unpaired *t*-test, (G-I) two-way ANOVA significant main effect of treatment, Bonferroni post hoc comparison. * p < 0.05, ** p < 0.01, *** p < 0.001, **** p < 0.0001. SA = spontaneous activity.

Patients with Fabry disease often suffer from mechanical allodynia in the hands and feet, and Fabry rodent models exhibit behavioral hypersensitivity to mechanical stimulation of the hindpaw [4, 13, 23]. Therefore, we next asked whether peripheral sensory nerves in Fabry rats exhibit altered responses to innocuous or noxious mechanical stimuli applied to the hindpaw. Single units recorded in dorsal roots from anesthetized Fabry and WT rats were classified as C-, Aδ, or Aβ fibers based on conduction velocity as measured via electrical stimulation of the sciatic nerve. A graded, innocuous von Frey stimulation (6, 16, 29g) was applied to the plantar foot and unit response was recorded. All classes of fibers, including Aβ, Aδ, and C-units, from Fabry rats exhibited increased firing frequency in response to innocuous mechanical stimulation of the hindpaw (Figure 1F-I). Furthermore, all fiber subtypes recorded in Fabry dorsal root nerves exhibited increased firing frequency in response to noxious mechanical stimulation (modified von Frey filament with a finely pointed tungsten tip) (Figure 1 J-M). These data indicate that sensory neurons in Fabry rats exhibit spontaneous activity and enhanced responses to mechanical stimuli, consistent with a large and small fiber sensory neuropathy.

An established source of spontaneous activity such as we recorded is the regenerative processes initiated in the growth cone of injured axons during their regrowth [36]. We have previously examined fiber counts in the peripheral nerves of Fabry rats and found decreased sensory fiber density of myelinated and unmyelinated subtypes compared to controls, as well as glycosphingolipid accumulation within axons [37]. Additionally, our data suggest that Fabry DRGs fail to show a change in the total number of neuron somata and there is an absence of Nageotte nodules, or satellite glial cell proliferation that follows degeneration of somata [38], which would be expected with loss of sensory somata (Figure 1S A, B). Together, these data support the interpretation that glycosphingolipid accumulation in Fabry disease causes denervation, which leads to established events [39–41] that result in spontaneous activity and could contribute to pain behaviors.

### Small diameter sensory neuron somata from Fabry rats exhibit spontaneous activity and hyperexcitability

The sensory neuron soma is the established model for assessing excitability of the sensory neuron membrane. Accordingly, we examined excitability of dissociated sensory neuron somata from Fabry rats. We observed that DRG somata from Fabry rats retain accumulated substrate after dissociation as we have previously published [23]. Figure 2A shows that a higher proportion of small diameter DRG neuron somata from Fabry rats display spontaneous activity compared to WT control. Fabry neuron somata exhibited reduced action potential current thresholds (rheobase), or the minimal current required to elicit an action potential (Figure 2B). Fabry neurons also showed increased firing frequencies evoked by suprathreshold depolarizing current injections compared to WT somata (Figure 2C-D). Other passive and current-evoked membrane properties recorded from small diameter DRG neurons were not different between genotypes (Table 1). Neuron somata larger than 32μm exhibited no differences to any excitability parameters as shown in Figure 2S and Table 2. Although dissociation may alter the functional properties of the soma, our observations support the possibility that hyperactivity of small diameter nociceptor somata may also contribute to the spontaneous activity of single units recorded in the dorsal root and pain behaviors observed in Fabry rats.

**Figure 2:**
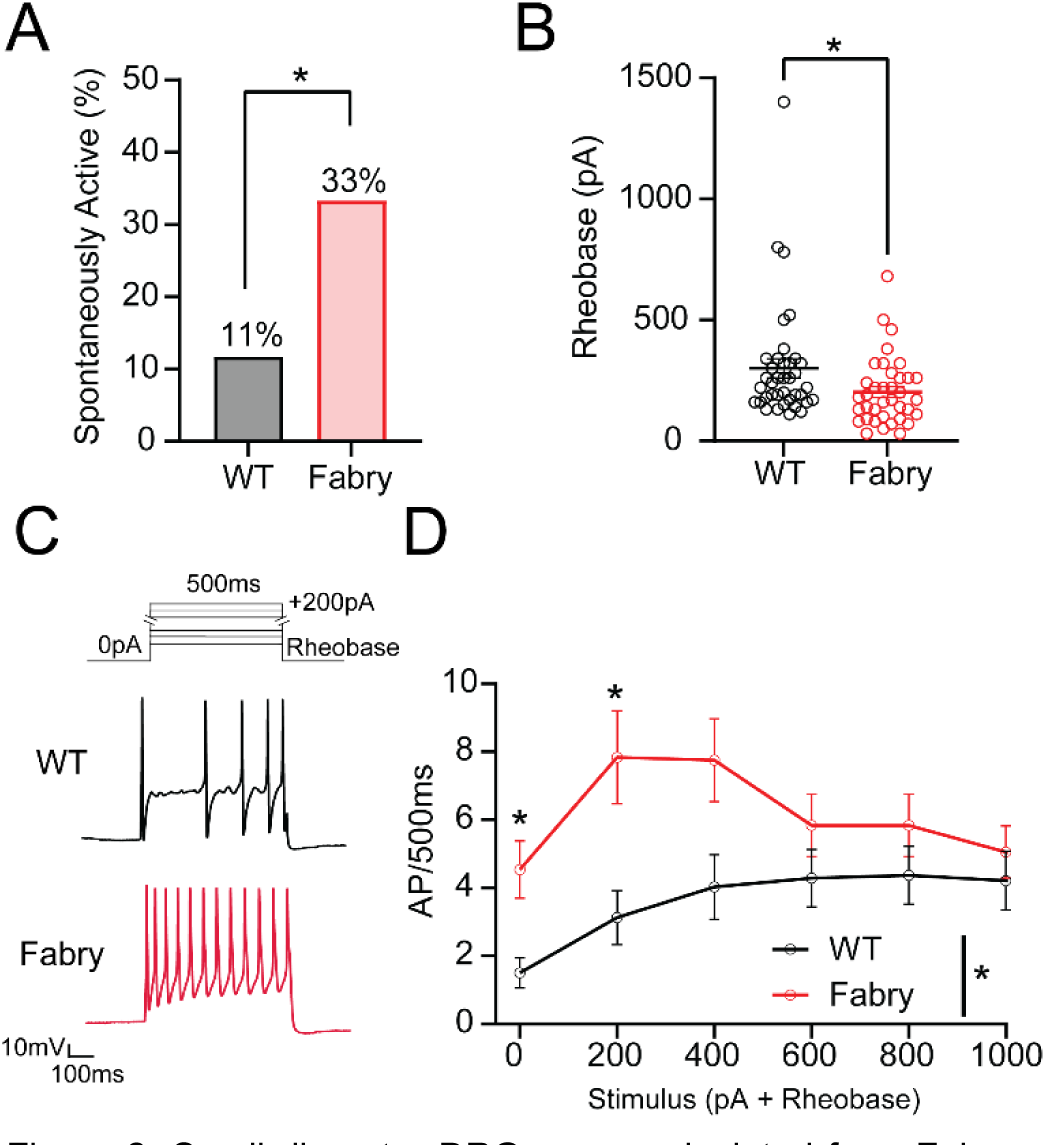
Small diameter DRG neurons isolated from Fabry rats exhibit spontaneous activity and current-evoked hyper-excitability. A) A higher proportion of isolated Fabry DRG neuron somata exhibit spontaneous activity compared to WT. B) Minimal action potential current threshold (rheobase) of Fabry neurons is reduced compared to WT neurons. C) Current protocol used for determining current-evoked firing frequency of Fabry and WT neurons with representative traces of Fabry and WT neurons undergoing current stimulation of 200 pA above rheobase for 500 ms. D) Fabry DRG neurons exhibit an increased firing frequency due to suprathreshold current stimulation. (A) n = 33-34 neurons from 6 animals per genotype; (B-D) n = 37-38 neurons from 8 animals per genotype. Data for (A) reported as mean, (B, D) as mean ± SEM. (A) χ^2^ Fisher’s exact post hoc comparison, (B) unpaired *t*-test, D) two-way repeated measures ANOVA significant main effect of treatment, Bonferroni post hoc comparison. * p < 0.05. RMP = resting membrane potential, AP = action potential.

### Schwann cells in Fabry rats exhibit abnormal morphology

Next, we asked what may drive the hyperexcitability we observed in the sensory neuron membrane. Numerous observations indicate extensive signaling between sensory neurons and their surrounding glia ([42] for review) and clinical tissues from patients with Fabry disease show morphological changes to Schwann cells [11, 13, 18], which suggests that glial signaling may disrupt the function of sensory neurons. We first examined whether Fabry rats exhibit abnormal morphology of Schwann cells by measuring the myelin sheath around neurons within crosssections of the tibial nerve using g-ratio analysis (ratio of the inner axonal diameter to total outer diameter) (Figure 3A). This indicated that axons in Fabry nerves have thicker myelin sheaths than WT neurons (Figure 3B; g-ratio plotted per axon diameter in Figure 3S). Peripheral nerves from patients with Fabry disease show evidence of denervated non-myelinating Schwann cells, which migrate and form regenerative tracks called Büngner bands to facilitate directional axon growth through release of growth factor proteins, and are a key characteristic of nerves undergoing Wallerian degeneration [43–45]. To assess this, we examined saphenous nerves from Fabry rats by electron microscopy which revealed denervated non-myelinating Schwann cells (Figure 3C) and evidence of regenerating axon tracks within or near degenerated myelinated axons [46] (Figure 3D). These morphological changes support our hypothesis that the interrelation between Schwann cells and peripheral sensory neurons is disrupted in Fabry disease.

**Figure 3:**
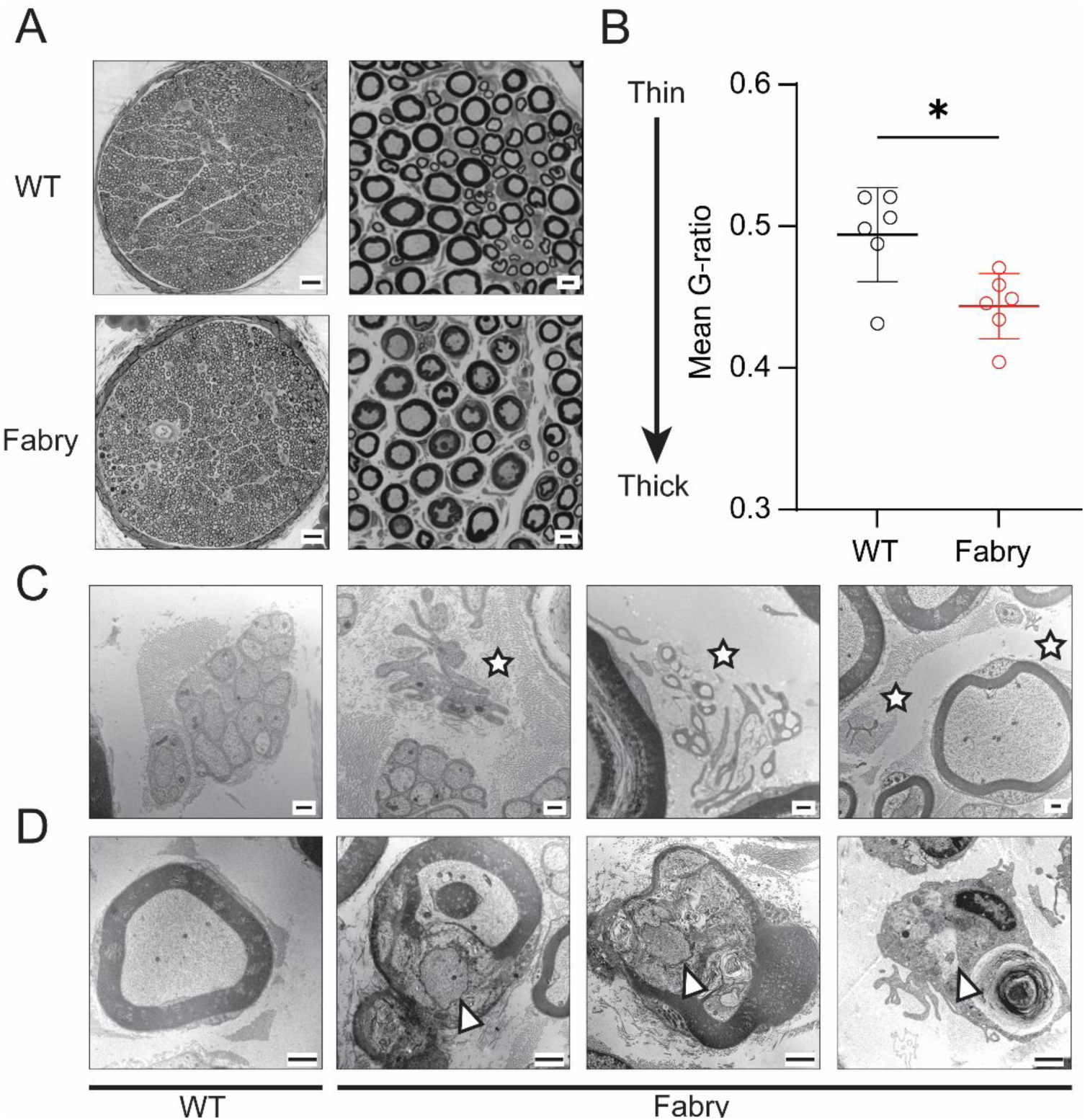
Schwann cells in Fabry sensory nerves are disrupted morphologically. A) Left: Representative light microscopy images of tibial nerve from Fabry and WT rats, scale bar 50μm; Right: Magnified images of peripheral nerve used to assess myelin architecture, scale bar 5μm. B) Analysis of mean G-ratio suggests that myelin sheath surrounding axons in Fabry nerves is thicker than myelin sheath surrounding WT axon. C) Representative transmission electron microscopy images of denervated Schwann cells (*), or Büngner bands, in the Fabry peripheral nerve, but not in the WT nerve, scale bar 0.5 μm. D) Representative transmission electron microscopy images show the presence of regenerating axons within or near degenerating myelinated fibers in the Fabry saphenous nerve (^) but not in the WT nerve, scale bar 1 μm. (A-B), tissue from n = 6 animals per genotype, one tibial nerve fascicle analyzed per animal; (C-D) tissue from n = 4 animals per genotype. (B) reported as mean ± SEM plotted per individual animal fascicle, unpaired *t*-test, * p < 0.05.

### Fabry Schwann cell mediators sensitize rat sensory neuron somata

Since Schwann cells in other neuropathic injury models release paracrine signaling compounds that alter ion channel function of sensory neurons [47–49], we further hypothesized that intrinsic hyperexcitability of the Fabry sensory neuron membrane may be induced by mediators released from Schwann cells. To test this, we cultured nearly pure (>90%), viable Schwann cells isolated from sciatic nerves of Fabry or WT rats [50] (Figure 4S A-C) and collected Schwann cell conditioned media (SCM) 3 days after plating. Both Fabry and WT Schwann cells exhibited robust and concentration-dependent calcium influx in response to ATP, suggesting Schwann cells from both cohorts are healthy (Figure 4S D, E). Naïve small diameter DRG neuron somata were then treated with SCM from Fabry, WT, or unconditioned (CTRL) groups overnight, which was washed out with extracellular recording solution. Within one hour, excitability was measured using current-clamp recordings. Sensory neurons treated with Fabry SCM displayed significantly depolarized resting membrane potentials (RMP) compared to neurons treated with either WT Schwann cell media or exposed to unconditioned media (CTRL) (Figure 4B). Because an elevated resting membrane potential facilitates spontaneous firing, we next evaluated non-evoked spontaneous activity. We found that a higher proportion of sensory neurons exposed to Fabry SCM exhibited spontaneous firing (Figure 4C-D). Naïve neurons also exhibited increased current-evoked firing frequency following Fabry SCM treatment (Figure 4E-F). Other passive or active membrane properties relevant to excitability, such as rheobase and action potential characteristics, were not different between treatment groups (Table 3). These data suggest that mediators secreted from Fabry Schwann cells may contribute to the development of hyperexcitability observed in Fabry sensory neurons.

**Figure 4:**
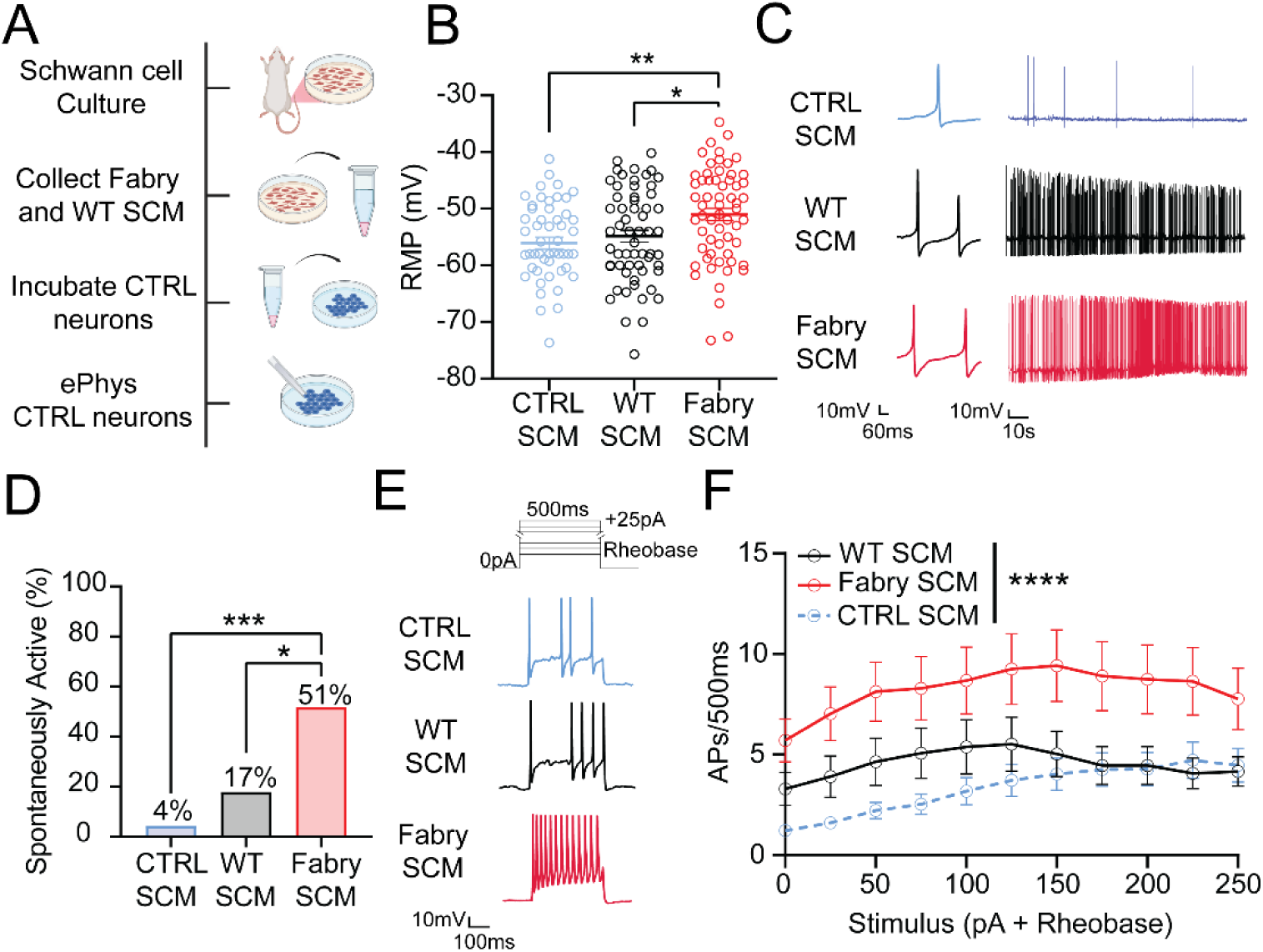
Fabry Schwann cell mediators induce peripheral neuron sensitization. A) Cultured DRG neuron somata from naïve Sprague Dawley (CTRL) rats were incubated overnight with either unconditioned CTRL, WT, or Fabry SCM. B) Neurons exposed to Fabry SCM demonstrated significantly depolarized resting membrane potentials. C) Representative traces of spontaneous firing from neurons incubated with CTRL, WT, and Fabry SCM over two minutes. D) Incubation of Fabry SCM caused more neurons to exhibit spontaneous activity. Neurons that fired one or more action potentials at RMP were considered spontaneously active. E) Current protocol and representative current-evoked traces from neurons exposed to CTRL, WT, or Fabry SCM undergoing stimulation of 150 pA above rheobase for 500 ms. F) Fabry SCM enhanced the firing frequency of neurons compared to neurons exposed to WT or CTRL SCM. WT and Fabry SCM derived from n = 5 individual animal Schwann cell cultures per genotype. (B) n = 51-60 neurons per treatment from 21 animals; (C-D) n = 23-29 neurons per treatment from 8 animals; (E-F) n = 28-31 neurons per treatment from 13 animals. Data reported as (B, F) mean ± SEM; (D) mean. (B) one-way ANOVA with Bonferroni post hoc comparison and (F) twoway repeated measures ANOVA with main effect of treatment (*** p < 0.0001). (D) χ^2^ with corrected Fisher’s exact post hoc comparison. * p < 0.05, ** p < 0.01, *** p < 0.001. RMP = resting membrane potential, SCM = Schwann cell conditioned media. AP = action potential.

### Fabry Schwann cell mediators evoke peripheral mechanical hypersensitivity

Hyperexcitability and spontaneous activity of nociceptors is associated with behavioral pain [51–53]. We therefore examined whether mediators released from Fabry Schwann cells may directly sensitize peripheral nerve terminals *in-vivo* and induce mechanical hypersensitivity, which is often associated with evoked pain in humans [54]. To test this, we injected Fabry or WT SCM into the glabrous skin hindpaw of naïve rats and assessed behavioral responses to mechanical stimulation (Figure 5A). We found that rats injected with Fabry SCM displayed robust mechanical hypersensitivity from 1 through 24 hrs post-injection, as assessed by von Frey stimulation (Figure 5B). We also asked whether animals exposed to Fabry SCM exhibit mechanical hyperalgesia to noxious needle stimulation, however, we observed no differences between treatments (Figure 4C). These data support the view that Fabry Schwann cells release mediators that sensitize peripheral nerve terminals.

**Figure 5:**
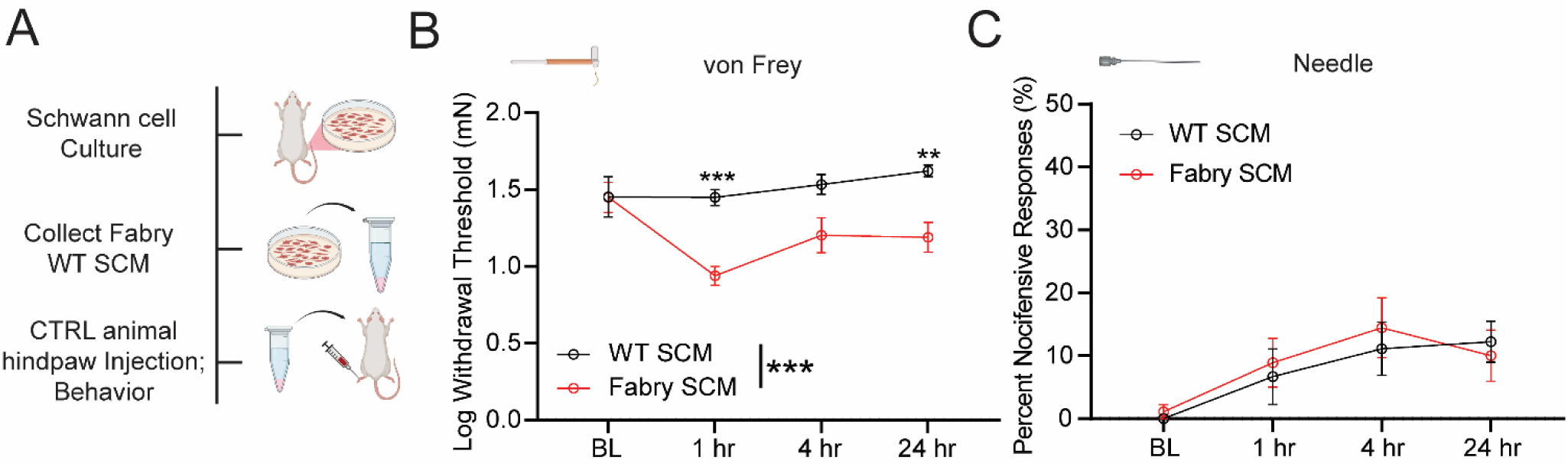
Naïve rats exhibit mechanical hypersensitivity due to peripheral injection of Fabry SCM. A) Naïve Sprague Dawley rats were given intraplantar injections of WT or Fabry SCM and underwent von Frey and needle poke stimulation of the hindpaw. B) Naïve rats injected with Fabry SCM exhibited increased mechanical hypersensitivity based on reduced 50% von Frey withdrawal thresholds. C) Rats exposed to either WT or Fabry SCM displayed a similar nocifensive response frequency to needle poke. (B,C) n = 9 animals per treatment, reported as mean ± SEM, two-way repeated measures ANOVA main effect of treatment (*** p < 0.001) with Bonferroni post hoc comparison. ** p < 0.01, *** p < 0.001. BL = pre-injection baseline behavioral measurements, SCM = Schwann cell conditioned media.

### Fabry Schwann cells release the protein p11 which induces mechanical hypersensitivity and sensitizes sensory neurons in naïve rats

We next discerned which compounds released from Fabry Schwann cells may be driving sensory neuron hyperexcitability. Schwann cells upregulate protein secretion during states of nerve injury [55–57], which we hypothesize may drive sensitization of sensory neurons. Therefore, we characterized the composition of Fabry and WT SCM using unbiased proteomic mass spectrometry. We identified 339 proteins in Fabry and WT SCM. Of the identified proteins, Fabry SCM showed upregulation of four proteins and downregulation of seven proteins compared to WT SCM (Table 4). One of the most highly upregulated proteins in Fabry SCM was the protein p11, also known as S100A10 (Fig 6A). Prior literature has shown that deletion of sensory neuron p11 reduces pain-like behaviors in a spinal nerve ligation model of neuropathic pain [58]. However, it is not known whether p11 induces sensory neuron hyperexcitability.

**Figure 6:**
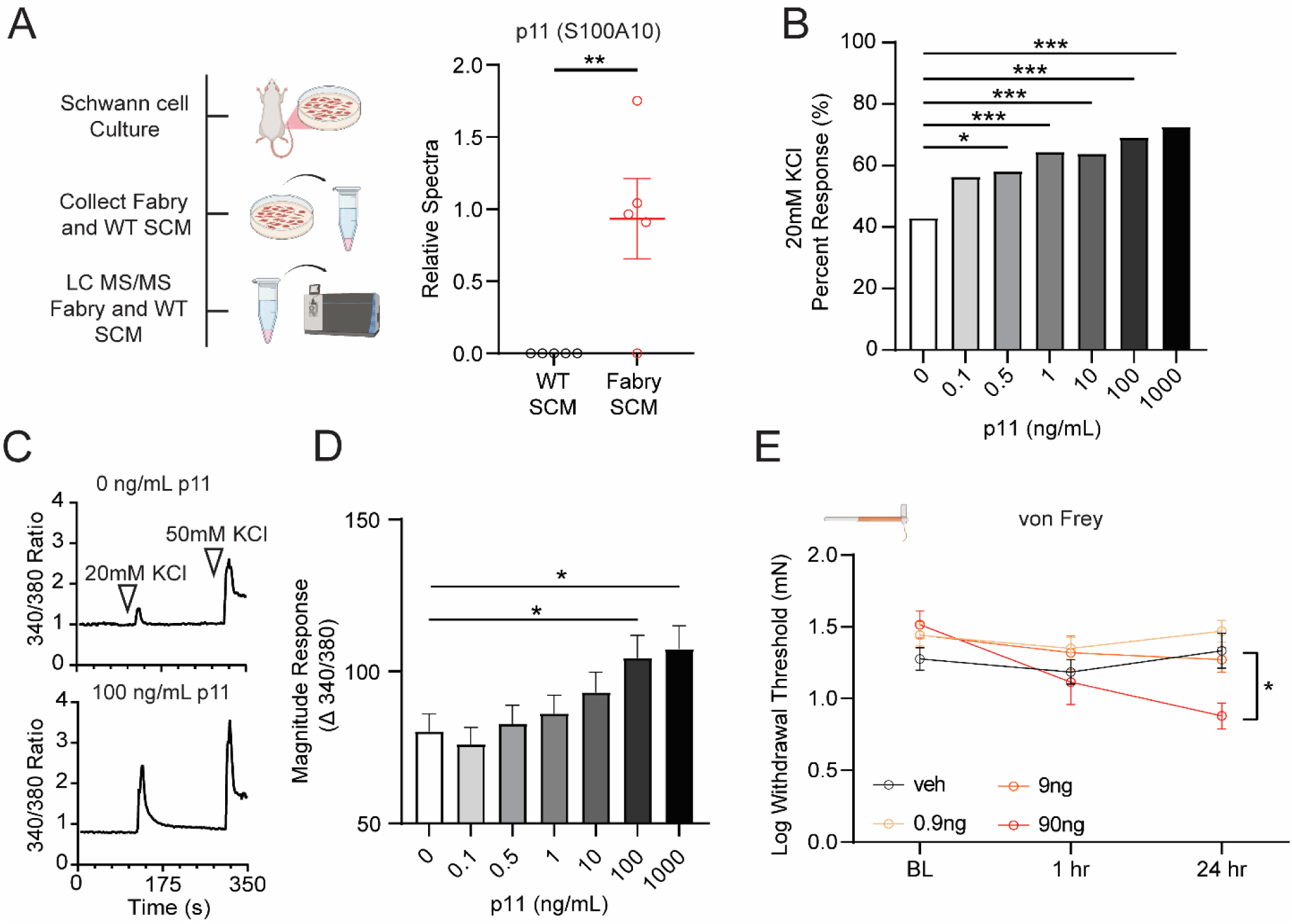
Fabry Schwann cells release the protein p11 (S100A10) which alters isolated DRG neuron function and induces pain-like behaviors in rats. A) Fabry and WT SCM were collected and underwent nanoLC MS/MS analysis, Fabry SCM contained significantly more p11 (S100-A10) compared to WT samples, refer to Table 4 for a list of the other significantly altered proteins. B) Significantly more neurons incubated with soluble p11 at various concentrations (0.5-1000 ng/mL) responded to mild depolarization with 20mM KCl as assessed using calcium imaging. C) Representative calcium imaging traces of 0ng/mL and 100 ng/mL p11 incubated neurons exposed to 20mM KCl for 10s, 50mM KCl used as a positive control. D) Neurons incubated with soluble p11 exhibited increased 20mM KCl-induced calcium flux. E) Intraplantar injection of p11 into naïve Sprague Dawley rats decreased hindpaw withdrawal thresholds to von Frey stimulation. (A) n = SCM samples from 5 animals per genotype, plotted per animal; (B-D) n = 140-160 neurons per treatment from 3 animals; (E) n = 8 animals per dose. (A, D-E) reported as mean ± SEM, (B) reported as mean. (A) Benjamini-Hochberg corrected two-tailed *t*-test, (B) χ^2^ with corrected Fisher’s exact post hoc comparison, (D) one-way ANOVA with Bonferroni post hoc comparison, (E) two-way repeated measures ANOVA with Bonferroni post hoc comparison. * p < 0.05, ** p < 0.01, *** p < 0.001. SCM = Schwann cell conditioned media.

To test whether p11 can directly activate or sensitize neurons, we incubated various concentrations of recombinant p11 onto naïve sensory neuron somata overnight with one hour washout and then exposed neurons to a depolarizing stimulus, during which we measured *in-vitro* calcium influx as an indirect correlate to neuronal activity [59]. Overnight incubation of 0.5-1,000ng/mL p11 increased the proportion of DRG neurons that responded to addition of 20mM KCl depolarization stimulus in the extracellular bath solution (Figure 6B), with incubation of 100-1,000ng/mL p11 also increasing response magnitudes (Figure 6C-D). We then investigated the effects of recombinant p11 on peripheral terminals *in vivo* through injection into the glabrous hindpaw of naïve rats and assessing mechanical thresholds. Rats injected with p11 exhibited mechanical hypersensitivity (Figure 6E), establishing a concentration-dependent response in which p11 sensitizes peripheral nerve terminals and isolated sensory neuron somata.

### p11 contributes to sensory neuron hyperexcitability induced by Fabry Schwann cells

As p11 sensitizes sensory neurons to depolarizing stimuli, we then asked whether p11 directly contributes to Fabry SCM mediated neuronal hyperexcitability. We incubated naïve DRG neuron somata overnight with 100ng/mL recombinant p11 and thereafter performed current-clamp recordings after one hour washout. Neurons incubated with p11 displayed significantly depolarized resting membrane potentials (Figure 7A). Incubation of p11 also decreased the rheobase of naïve sensory neurons (Figure 7B). Furthermore, neurons fired more action potentials in response to depolarizing current injections following p11 incubation (Figure 7C). We found that p11 incubation induced a trending, but not significant, increase in the proportion of neurons exhibiting spontaneous activity (Figure 5S A-C), and that other membrane properties related to excitability remained similar between treatments (Table 5). Prior literature shows that deletion of neuronal p11 reduces voltage-gated sodium channel function in DRG neuron somata [58], which may influence excitability [60]. We tested whether p11 enhances voltage-gated sodium channel function through voltage-clamp recordings of naïve sensory neurons incubated overnight with p11 that was washed out prior to recording (Figure 7 E-G). Neurons exposed to p11 exhibited increased voltage-dependent sodium channel current densities compared to control neurons (Figure 7 F-G).

**Figure 7:**
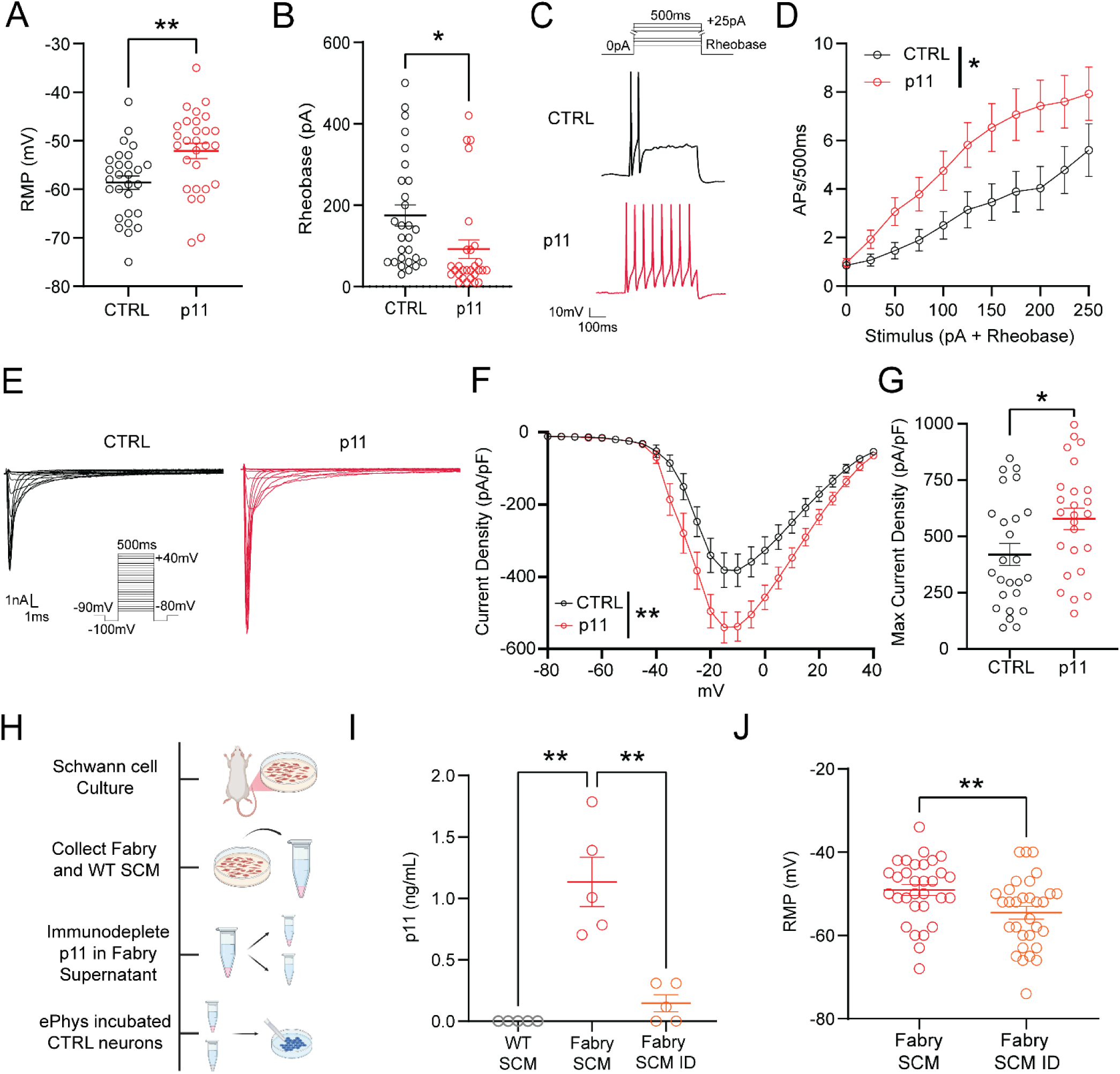
p11 contributes to the resting membrane depolarization of naïve DRG neurons treated with Fabry SCM. A) Naïve neurons exposed to 100 ng/mL p11 exhibited significantly depolarized resting membrane potentials compared to CTRL neurons. B) Incubation of p11 reduced the rheobase of naïve neurons. C) Current protocol and representative traces of neurons undergoing current stimulation of 200 pA above rheobase for 500 ms. D) Neurons exposed to p11 exhibit increased firing frequency to suprathreshold current stimulation compared to CTRL. E) Voltage protocol and representative current traces from neurons incubated with 100 ng/mL p11 or CTRL media that underwent analysis of sodium channel currents (representative traces depict currents elicited from -60mV to +5mV). F) Incubation with 100ng/mL p11 enhanced peak sodium current densities. G) The maximum inward current density was higher in neurons exposed to p11, reported as absolute values. Mean capacitance values (pF ± SEM) for neurons tested were 25.5 ± 2.0 and 23.5 ± 1.9 for CTRL and p11 incubated neurons respectively. H) Fabry SCM was collected and underwent p11 immunodepletion with control Fabry SCM media and placed onto naïve neurons. I) Immunodepletion reduced p11 concentration of Fabry SCM by 86%, with WT SCM exhibiting concentrations below the limit of detection as measured by ELISA. J) Naïve neurons exposed to Fabry SCM ID exhibited a hyperpolarized RMP compared to neurons treated with Fabry SCM. Values reported as mean ± SEM. (A-D) n = 28 neurons per treatment from 7 animals; (E-G) n = 26 neurons per treatment from 8 animals; (I) n = 5 WT and Fabry Schwann cell cultures from individual animals; (J) n = 31 neurons per treatment from 8 animals. (A, B, G, J) unpaired *t*-test, (D, F) two-way repeated measures ANOVA with main effect of treatment, (I) two-way ANOVA, Bonferroni post hoc comparison. * p < 0.05, ** p < 0.01. RMP = resting membrane potential, SCM = Schwann cell conditioned media, SCM ID = Schwann cell conditioned media with immunodepleted p11, AP = action potential.

We then investigated whether p11 is a critical compound released from Fabry Schwann cells to induce sensory neuron hyperexcitability. To test this, we collected Fabry Schwann cell conditioned media and immunodepleted p11 using dynabead-conjugated anti-p11 antibody, which was removed through magnetic sorting (Figure 7H). Removal was confirmed through a commercially available enzyme-linked immunosorbent assay (ELISA) for p11. Fabry SCM contained an average of 1.13 ng/mL p11 before immunodepletion (Figure 7I). After immunodepletion, the level of p11 was reduced by 86% (Figure 7I). The concentration of p11 in the immunodepleted SCM (mean 0.15 ng/mL) was below the effective concentration in which p11 induced significant alteration to neuronal activity as shown in Figure 6C-D. Therefore, we incubated naïve neuron somata with standard Fabry SCM or Fabry SCM with immunodepleted p11 (Fabry SCM ID) and used current-clamp electrophysiology to detect differences in spontaneous activity or excitability. Neurons incubated with Fabry SCM ID exhibited a significantly hyperpolarized resting membrane potential relative to standard Fabry SCM (Figure 7J). However, removal of p11 did not influence the proportion of neurons that exhibited spontaneous firing, neuron rheobase, current-evoked firing frequency (Figure 6S A-C), or other passive and current-evoked membrane properties related to excitability compared to Fabry SCM (Table 6). These findings suggest that release of p11 from Fabry Schwann cells contributes to the hyperexcitability of sensory neurons through its influence on resting membrane depolarization, thereby possibly contributing to pain in Fabry disease.

## Discussion

Chronic pain that afflicts patients with Fabry disease is critically understudied and its underlying causes have received limited research attention. Using a genetic rat model of Fabry disease, we discovered that peripheral sensory neurons in Fabry rats display prominent spontaneous activity and hyperexcitability. Because Schwann cells in the nerves of Fabry patients and in our rat model are disrupted morphologically, we investigated whether Schwann cells may contribute to the pathological hyperexcitability in Fabry sensory neurons. This revealed that Fabry Schwann cells release factors that induce hyperexcitability in naïve DRG neurons and induce behavioral mechanical hypersensitivity in naïve rats. The release of the candidate algogen p11 (S100A10) from Fabry Schwann cells is upregulated, and recombinant p11 induces hyperexcitability in naïve sensory neurons. Removal of p11 from Fabry Schwann cell media causes relative hyperpolarization of neuronal resting membrane potential, indicating that release of p11 contributes to the excessive neuronal excitability caused by Fabry Schwann cells. Together, our findings suggest that communication between Schwann cells and sensory neurons in the peripheral nerve may drive enhanced nociceptive signaling seen in Fabry disease.

Patients with Fabry disease experience a variety of pain types, including debilitating shooting or burning pain in the hands and feet that presents in childhood onward [4, 61, 62], but our knowledge of what drives this pain is limited. This characteristic pain phenotype may be caused by increased nociceptor excitability [63, 64], but testing this fundamental hypothesis has been difficult due to the lack of appropriate models that recapitulate hypersensitivity to painful stimuli as experienced by patients. Here, we propose that the rat model appropriately replicates the mechanical hypersensitivity and ongoing pain seen in patients with Fabry disease, and that this pain is due to underlying nociceptor hyperexcitability. First, we have previously shown that the Fabry model rat exhibits mechanically evoked-pain [23] as well as spontaneous pain [24] phenotypes similar to patients. In this present study, we demonstrate that individual sensory neurons have significant spontaneous activity and increased sensitivity to mechanical stimuli of all sensory fiber types in Fabry rats, which establishes that the peripheral portion of sensory neurons are a critical site of pathologic function in Fabry disease. Isolated small diameter sensory neuron somata exhibit current-evoked hyperexcitability, in addition to being hypersensitive to mechanical stimuli as we previously showed [23], thereby indicating that increased intrinsic sensory neuron membrane excitability underlies mechanical sensitivity and spontaneous neuronal activity seen in Fabry rats.

Further, this model replicates Schwann cell denervation as shown here, as well as sensory fiber loss as we previously shown [37]. Together, these results show that the Fabry rat model is suitable for studying the complex mechanisms that drive this debilitating yet understudied genetic pain disease.

Emerging evidence suggests that Schwann cell dysfunction plays a role in the development and maintenance of chronic pain. Opto- or chemogenetic activation of Schwann cells induces peripheral sensory nerve sensitization and pain-like behaviors in rodents [19–22], and Schwann cells can release specific algogens that can cause behavioral pain in rodents [47–49, 65]. Here, we identify a direct link by which Schwann cells induce nociceptive neuron hyperexcitability through mediator release and provide an unbiased characterization of these putative algogens. We found that sensory neuron somata from naïve rats exposed to Fabry Schwann cell mediators developed hyperexcitability, including increased spontaneous activity and current-evoked firing frequency. Using intact animals, injection of Fabry Schwann cell mediators into the hindpaw induced mechanical hypersensitivity, showing that these mediators can activate the peripheral processes of sensory neurons. Additionally, we have shown that Fabry Schwann cells exhibit morphological pathology near peripheral sensory axons *in-vivo*. Prior evidence suggests that Schwann cells can induce gene expression changes in neurons [66] and alter ion channel organization and function in adjacent axons and neurons that are relevant for neuron excitability, such as voltagegated sodium channels [67–70]. We think it is highly likely that Schwann cells contribute to enhanced excitability of Fabry sensory neurons by secreting mediators that alter neuronal gene and ion channel function.

Our study demonstrates that Fabry Schwann cells upregulate the release of the protein p11 (S100A10). We further show that p11 exerts previously uncharacterized regulation of nociceptive signaling, including enhanced sensory neuron excitability and induction of mechanical hypersensitivity upon peripheral hindpaw injection into naïve rats. It is unknown how p11 induces sensory neuron hyperexcitability. Little is known about p11 as a putative paracrine signaling compound, but this small (11 kDa) protein is present in many cell types and is associated with exosomal secretion pathways [71], which is a potential mechanism for release of p11 from Fabry Schwann cells and subsequent DRG neuron internalization. Intracellular p11 modulates the trafficking and subsequent localization of many ion channels to and from cellular membranes in neurons that are relevant for excitability, which has been previously reviewed [72]. This includes the voltage-gated sodium channel Nav1.8 [58, 73, 74]. Our findings show that exogenously applied p11 enhances voltage-gated sodium ion channel currents (Figure 7 E-G), aligning with a prior study suggesting that knockdown of p11 within neurons can reduce these currents [58]. Additionally, we show that incubation of p11 enhances the calcium influx of naïve neurons exposed to KCl (Figure 6 B-C) which may be due to increased membrane localization of L-type calcium channels [75] by p11 [76]. Removal of p11 from the Fabry SCM causes a selective hyperpolarization of the resting membrane potential compared to neurons treated with complete Fabry SCM (Figure 7G, Figure 6S, Table 6). This finding suggests that elevated p11 depolarizes the resting membrane potential of neurons by increasing hyperpolarization-activated cyclic nucleotide-gated (HCN) or decreasing two-pore domain (K2P) TASK1 membrane localization [77], as prior literature shows that knockdown of p11 in neurons reduces HCN channel-mediated currents [78] and increases putative TASK1 currents [79]. Considering this evidence, we propose that Fabry Schwann cells secrete p11 to modify the localization of various ion channels to increase the excitability of sensory neurons.

We recognize the limitations of our study. Although we have shown that mediators from Fabry Schwann cells, including p11, contribute to mechanical hypersensitivity and peripheral sensory neuron dysfunction observed in Fabry rats, we have not determined if Schwann cells drive these observations *in vivo*. This issue is difficult to address as the cellular mechanisms through which Fabry Schwann cells release algogenic mediators are unknown, including release of p11. Systemic knockdown of p11 would induce off-target effects [80], and no techniques are available to functionally neutralize p11 released from Schwann cells. Further investigation of the cellular machinery required for Schwann cell secretion is needed. In addition, while p11 contributes to the resting membrane potential depolarization observed in sensory neurons exposed to mediators from Fabry Schwann cells, mediators that increase spontaneous activity or other aspects of hyperexcitability, including decreased rheobase, in these treated sensory neurons remain unexplained. Finally, further characterization of the Fabry Schwann cell secretome is necessary to uncover how interaction between Schwann cells and neurons drive nociceptor sensitization and chronic pain in Fabry disease.

We have shown that peripheral nerve dysfunction and the associated hyperexcitability of sensory neurons underlies chronic pain in this Fabry disease model, and that sensory neuron hyperexcitability may result from chronic altered signaling by Schwann cells. These findings indicate that similar processes are potential mechanisms for the heightened nociceptive signaling seen in this understudied disease. Further study on peripheral sensory neuron and glial dysfunction in Fabry disease may uncover novel therapeutic targets for this debilitating genetic pain disorder.

## Methods

### Animal model

The X-linked genetic Fabry disease (Fabry) rat model [23] (Rat Genome Database symbol: Glaem2Mcwi) was used for *in vivo* dorsal root teased single unit recordings and for collecting primary sensory neuron and Schwann cell cultures, compared to age and sex-matched wild-type (WT) littermate controls. All rats used for in-vivo and in-vitro electrophysiology and collection of Fabry Schwann cells were hemizygote males between 20-40 weeks old. For evaluating the effects of Schwann cell media or p11 protein treatment on naïve sensory neuron function, male Sprague Dawley outbred rats (Envigo) aged 15-30 weeks were used (CTRL). For animal behavior experiments, both male and female Sprague Dawley rats were used. No differences related to sex were observed in behavior studies, so sexes were combined for analysis.

### Light microscopy (LM) and transmission electron microscopy (TEM)

#### Peripheral nerves

Nerves from Fabry and WT rats were collected [37] and processed [81, 82] as previously described. Briefly, nerves were fixed with 4% paraformaldehyde and 2% glutaraldehyde in sodium cocodylate buffer at 4°C overnight, then postfixed with 1% osmium tetroxide for 2 hours at room temperature. Samples were dehydrated in a series of methanol gradients and infused with epon. Sections 0.5 μm thick were cut for light microscopy analysis and stained with toluidine blue, while thin sections of 70 nm were prepared for transmission electron microscopy (TEM), stained with 25% uranyl acetate and counterstained with lead citrate. Cross-sections of the tibial nerve were examined using light microscopy to manually assess myelination through G-ratio analysis (quantification of inner axonal diameter to total outer diameter). Axons within the entire nerve fascicle were counted per animal and analyzed using ImageJ software. TEM cross-sections of the saphenous nerve were obtained using a JEOL 1400 Flash electron microscope operating at 80KV and assessed for morphological abnormalities. G-ratio calculation and TEM image acquisition were performed by researchers blinded to genotype.

#### Dorsal root ganglia (DRG)

Tissue from Fabry and WT rats aged 13 weeks were harvested and processed as previously described [23, 83]. DRG were fixed with 10% neutral-buffered formalin and then paraffin-embedded. Blocks were cross-sectioned 4 µm thick, deparaffinized, and then stained with hematoxylin and eosin (H&E). Slides were imaged using a Hamamatsu (Hamamatsu, Japan) NanoZoomer 2.0-HT slide scanner. Neuron somata from the entire DRG cross-section were counted compared to the cross-sectional area of the DRG, and DRG were assessed for the presence of Nageotte nodules, or residual clusters of satellite glial cells associated with sensory neuron somata degeneration [38]. All analyzes were performed by researchers blinded to genotype.

### Cell culture

#### Sensory neuron somata

Dorsal root ganglia (DRGs) were harvested and neurons were dissociated and cultured as done previously [23, 84]. Specifically, rats were anesthetized (4% isoflurane), sacrificed, and lumbar DRGs were isolated. DRGs were incubated with 1mg/ml collagenase type IV (Sigma) for 45 min then 0.05% trypsin (Sigma) for 45 min. Neuronal somata were mechanically dissociated and plated onto laminin-coated coverslips. One hour after plating, neurons were fed cell media [high glucose DMEM, 2 mmol/L L-glutamine, 1% glucose, 100 units/mL penicillin, 100 ug/mL streptomycin, 2% heat-inactivated horse serum]. Neurons grew overnight at 37°C and 5% CO2. For experiments mentioned in the text, neurons were fed with Schwann cell collection media [high glucose DMEM (ThermoFisher), 10 nM neuregulin (Recombinant Heregulin-β1177–244, PreproTech), and 2 μM forskolin (Sigma)] that was extracted from Fabry or WT Schwann cell cultures. For some experiments, the inclusion of an unconditioned media control (CTRL) was incubated onto neurons overnight (12-16 hours), which was unconditioned Schwann cell collection media.

#### Schwann cells

Schwann cells were cultured using a previously published protocol [50] with modifications. Briefly, adult Fabry or WT rats were anesthetized, culled, and sciatic nerves were removed by transection under the gastrocnemius muscle and at the nerve trifurcation. The epineurium was removed, and nerve fascicles teased. These were subjected to enzymatic dissociation with 0.25% Dispase I (ThermoFisher) and 0.05% type I collagenase (EMD Milipore)) in high-glucose DMEM (ThermoFisher) with antibiotics overnight at 37 °C and 5% CO2.

Enzymatic dissociation was stopped with the addition of 40% fetal bovine serum, centrifuged, and cell pellets were resuspended in DMEM with 10% serum. Resuspended cells were mechanically dissociated, then plated onto air-dried PDL (ThermoFisher) and laminin-coated plates (Sigma). Two to three hours after plating, Schwann cells were washed with PBS to remove excess debris, and then fed with Schwann cell growth media [high glucose DMEM (ThermoFisher),10 nM neuregulin (Recombinant Heregulin-β1177–244, PreproTech), and 2 μM forskolin (Sigma), 10% FBS (ThermoFisher)]. To assess Schwann cell viability, cells were un-adhered with 0.05% trypsin then diluted 1:1 in trypan blue (Sigma) and viability was automatically counted (Countess 3, Invitrogen).

#### Schwann cell culture purity

Schwann cells were cultured for 2 days and then fixed in 4% PFA for 20 minutes, washed with PBS, then permeabilized with blocking solution (0.05 M Tris-buffered saline with 0.3% Triton-X) for 1 hr then washed. Recombinant anti-SOX10 antibody (Abcam, EPR4007) in blocking solution was added for 1 hr (1:500), then washed with PBS and secondary antibody in blocking solution (Alexa Fluor™ 594 donkey anti-rabbit IgG, ThermoFisher, R37119) was added (1:1000) for 1 h. After washing, coverslips were mounted with mounting media + DAPI (Vector Laboratories) and imaged using a Leica SP8 Upright Confocal Microscope. The middle of the coverslip was found, and one 4×4 stitched image (16 separate images) was taken from each culture at 20x magnification. Purity of the culture was quantified using a modified colocalization analysis we have previously described [37]. Briefly, 30 DAPI+ cells that were determined to be negative for SOX10 were manually assessed for mean SOX10 fluorescence. Any cell with SOX10 signal four standard deviations above this baseline mean were considered SOX10 positive; ratio of SOX10+ and DAPI+ cells compared to all DAPI+ cells was automatically calculated through an ImageJ macro and reported as percent purity. Researchers were blinded to genotype for analyses.

#### Schwann cell conditioned media (SCM)

Schwann cell cultures were grown for two days in Schwann cell growth media to 70-80% confluency. Cells were then washed with PBS and replaced with Schwann cell collection media [high glucose DMEM (ThermoFisher), 10 nM neuregulin (Recombinant Heregulin-β1177–244, PreproTech), and 2 μM forskolin (Sigma)] and cultured for 1-2 days. Schwann cells have been shown to grow in culture with or without crude serum [85]; serum was not included in collection media to reduce confounding variables for all experiments. This media was collected, filtered using a 0.22um filter to remove debris or cell fragments, flash-frozen in liquid nitrogen, and stored at -20C or -80C for less than one month for functional assays. Media samples, represented as either Schwann cell conditioned media or unconditioned media controls, were then thawed and used for all subsequent experiments. Media underwent a maximum of two flash-freeze cycles for all assessments and was not diluted.

#### p11 treatment

Recombinant rat protein p11 (S100A10 Recombinant Protein, Aviva Systems Biology, OPCD06771) was dissolved in distilled water to obtain a final concentration of 100ug/mL and stored at -80C for less than one month. Protein was further diluted in DRG neuron culture media to obtain the respective concentrations (0.1-1,000 ng/mL) for calcium imaging experiments and whole-cell patch clamp electrophysiology experiments; neurons were incubated with this protein overnight. For intraplantar injection to assess rodent behavior, protein was dissolved in saline to a dose of 0.9, 9, or 90 ng per 30uL injection, saline was used as vehicle (veh) injections.

#### p11 Immunodepletion

Fabry Schwann cell media was collected as described above. Media was split into two equal aliqouts: one undergoing immunodepletion and the other used as a control batch. Anti-p11 antibody (10ug) (S100A10 Polyclonal antibody, ProteinTech, 11250-1-AP) was incubated with 50uL Dynabeads® Protein A (Invitrogen) for 10 min, and supernatant removed with a DynaMag™-2 Magnet tube rack (Invitrogen). Afterwards, Fabry Schwann cell media was added the anti-p11 antibody bound Dynabeads® and let incubate on a rotating platform for 60 min at RT. Samples were then placed onto the magnetic tube rack and supernatant was removed. Depletion of p11 from the media was confirmed using Rat S100 Calcium Binding Protein A10 (S100A10) ELISA Kit (Biomatik, Cat# EKN48271-96T) as per the attached instructions.

### Dorsal root teased fiber single unit recording

Fabry and WT rats underwent *in vivo* dorsal root single unit recordings as we reported previously [86]. Briefly, rats were anesthetized with subcutaneous injection of urethane (100mg/kg) followed by isoflurane that was progressively reduced to 0.2% over 30 min. A laminectomy was performed to expose the spinal cord from the T13 to the L3 vertebrae and covered with warm mineral oil (36°C). Dura was removed, and rats were mounted on a spinal frame with stabilizing vertebral clamps. The L4 dorsal root was gently released from connective tissues and transected at the rootlets adjacent to the spinal cord. The dorsal root was placed onto a glass platform and teased into fine bundles that were individually placed onto a platinum/iridium recording electrode for observation of spontaneous and evoked activity. A reference electrode was placed in adjacent muscle tissue. Signals were collected with an Axoclamp 900 A microelectrode amplifier (Molecular Devices, San Jose, CA) and the headstage (HS-9A-x0.1U with feedback resistance of 100MΩ) was served as a preamplifier with gain setting of 500 or higher, filtered at 1 kHz, and sampled at 10 kHz using a digitizer (DigiData 1440 A, Molecular Devices). Action potentials were isolated by setting the threshold above background noise.

For recording spontaneous activity, bundles were observed for a 3 min observation period and recorded for 3–4 min if present. For recording evoked activity, the receptor field of a unit was identified by low intensity mechanical stimulation of the glabrous plantar skin of the hind paw with a small glass probe (with 1 mm round tip). Firing frequency to innocuous mechanical stimulation was then examined with graded von Frey monofilaments and a modified von Frey monofilament with a tungsten tip for noxious stimulation. To identify unit types, the sciatic nerve was stimulated and used to calculate conduction velocity by dividing the distance between stimulation and recording sites by the response latency of the electrically evoked action potential. Fiber types were then classified based on conduction velocity; 20m/s or greater for Aβ, between 3 and 20m/s for Aδ, and less than 3m/s for C-type. The experimenter was blinded to genotype for all analyses.

### Whole-cell patch clamp electrophysiology

Neuronal somata were categorized into either small (≤ 32µm) or larger diameter (> 32µm) as the electrophysiology properties of rodent neurons varies based upon size [87, 88], and the majority of small diameter neurons are considered nociceptors [89]. Neuronal capacitance was fully compensated and continuously monitored to ensure stable recording conditions. Whole-cell recordings were obtained using HEKA EPC10 amplifier, and recordings were obtained using Patchmaster Next software (version 1.2). The experimenter was blinded to genotype or treatment for all analyses. All reagents for patch clamp analysis were obtained from ThermoFisher unless otherwise specified.

#### Current-clamp recordings

Isolated sensory neuron somata were superfused with extracellular buffer (140 NaCl, 2.8 KCl, 2 CaCl2, 1 MgCl2, 10 HEPES, 10 glucose, and 8.8 sucrose, pH 7.4 ± 0.02 and 310 ± 3 mOsm, adjusted with sucrose). Borosilicate glass pipettes (2-6 MΩ) filled with internal solution (In mM: 135 KCl, 4.1 MgCl2, 2 EGTA, 0.2 GTP, 2.5 ATP, and 10 HEPES, pH 7.2 ± 0.02 and 290 ± 2 mOsm) were pulled using a Sutter Instruments P87 pipette puller and used to perform patch clamp recordings. Series resistance was maintained at <15 MΩ and compensated at 80%.

#### Spontaneous activity

Whole cell recordings were established in voltage-clamp mode, then switched to current-clamp mode to measure resting membrane potential (RMP). Voltage was recorded at RMP for 2 minutes to observe spontaneous, suprathreshold action potentials (>0mV) with zero current injection. Cells firing at least one action potential during the 2-minute period were considered spontaneously active.

#### Current-evoked excitability

Neuron somata were held at -70mV and intrinsic excitability was recorded using the following protocols [90]: (1) Voltage-current (V-I) relations were obtained using 20 sweeps of 500 ms alternating ascending/descending current pulses (5 pA steps from holding current). The plateau voltage deflection was plotted against current amplitude, and input resistance was determined from the slope of an IV plot. (2) Action potential (AP) properties were measured using an ascending series of 500 ms depolarizing current pulses. Rheobase was defined as the minimal current to elicit at least a single spike. AP threshold was determined from a derivative function, where dV/dt first exceeded 28 mV/ms. AP amplitude was determined relative to AP threshold, and AP half-width was measured as the width at half of the AP amplitude. (3) A series of eleven 500 ms depolarizing current steps (range, rheobase to 250 pA above rheobase; 25 pA increments, 20s intervals) or seven 500 ms depolarizing current steps (range, rheobase to 1,600 pA above rheobase; 200 pA increments, 20s intervals) were used to determine action potential firing frequency.

#### Voltage-clamp recordings

Extracellular buffer (In mM: 70 NaCl, 70 Choline-Cl, 3 KCl, 1 MgCl2, 1 CaCl2, 10 glucose, 10 HEPES, pH 7.35 ± 0.02 and 310 ± 3 mOsm, adjusted with sucrose) was flowed over isolated sensory neuron somata from control rats. The addition of 20 mM TEA-Cl and 0.1 mM CdCl2 was added to extracellular buffer to block voltage-gated K+ channels and Ca2+ channels respectively [91]. Fire-polished borosilicate glass pipettes (1-4 MΩ) were filled with internal solution (In mM: 140mM CsF, 10mM NaCl, 2mM MgCl2, 0.1 CaCl2, 1.1 EGDA 10 HEPES, pH 7.3 ± 0.02 and 310 ± 3 mOsm, adjusted with sucrose) and pulled using a Sutter Instruments P87 pipette. Somata were established in voltage clamp at a holding potential of -90mV. Series resistance was maintained at <10 MΩ and compensated on 85%, then held at holding potential for 2-4 min. Currents were elicited by incremental depolarizing steps (+5 mV increments, 500 ms duration, 5s intervals) between -80mV to +40mV, with a -100mV hyperpolarizing pulse given prior and after each step for 50ms. The average of three sweeps per neuron were taken for determination of peak current density at each step, which were filtered at 2.9 kHz.

### Calcium imaging

#### Schwann cells

Schwann cells were cultured in serum-containing media and plated onto glass coverslips. All reagents were obtained from ThermoFisher unless otherwise indicated. After two days in culture, coverslips were washed with extracellular buffer solution (150 mM NaCl, 10 mM HEPES, 8 mM glucose, 5.6 mM KCl, 2 mM CaCl2, and 1 mM MgCl2, pH 7.40 ± 0.03, and 320 ± 3 mOsm) for 30 minutes, incubated with 2.5 mg/mL Fura-2-AM (Life Technologies) then washed for 30 minutes. Experimenters were blinded to genotype, and coverslips were mounted onto a perfusion chamber, placed on an inverted fluorescent microscope, superfused with extra-cellular buffer at 6 mL/min, and imaged at 20x magnification. Under light microscopy, the experimenter marked 20-50 Schwann cell bodies per coverslip prior to imaging; all marked Schwann cells were included in final analysis. Fluorescence images were captured at 340 nm and 380 nm with a cooled Andor Zyla sCMOS camera (Oxford Instruments) to calculate the bound to unbound ratio (340/380). NIS Elements software (Nikon) was used to detect and analyze intracellular calcium changes. Differing concentrations of ATP (1, 10, 50 µM) were applied for 10 seconds to the bath. Schwann cells that exhibited a >20% 340/380 response magnitude compared to baseline within 30 s after ATP application were considered responders.

#### Sensory neuron somata

Calcium imaging of dissociated neuronal somata was conducted as we have previously published (Cowie et al., 2019) with the same equipment and reagents used for Schwann cell calcium imaging. Somata dissociated from DRGs were incubated overnight in media with or without 0.1-1000ng/mL p11. Somata were then incubated with Fura-2-AM for one hour and imaged. To induce membrane depolarization, we added 20mM KCl to the extracellular buffer solution, while reducing NaCl concentration to maintain an osmolarity of 320 mOsm. The depolarizing solution was applied to neurons for 10 seconds to determine both number of responding neurons and response magnitude. Somata that exhibited a ≥20% increase in 340/380 ratio within 30 seconds after KCl application compared to baseline ratio were considered positive responders. As a positive control, 50mM KCl was applied near the end of the recordings; somata were considered viable and subsequently analyzed only if they responded to 50mM KCl. The experimenter was blinded to genotype or treatment during all calcium imaging analyses.

### Mass spectrometry

#### Enzymatic “In Liquid” Digestion

Acellular rat cell culture media (∼4-5ml) were speed-vac to ∼250µl each, diluted 1:1 with water and proteins were extracted via TCA/Acetone precipitation for 45 minutes on ice to (10% TCA in 50% Acetone final vol:vol). Spun for 10 minutes at room temperature with max speed (16,000xg) and generated pellets were washed twice with cold (−20°C) acetone. Generated protein pellets were solubilized and denatured in 15μl of 8M Urea in 50mM NH4HCO3 (pH 8.5). Subsequently diluted to 60μl for reduction step with: 2.5μl of 25mM Dithiothreitol (DTT) and 42.5μl of 25mM NH4HCO3 (pH 8.5). Incubated at 52°C for 15 minutes, cooled on ice to room temperature then 3μl of 55mM Chloroacetamide (CAA) was added for alkylation and incubated in darkness at room temperature for 15 minutes. Reaction was quenched by adding 8μl of 25mM DTT. Finally, 8μl of Trypsin/LysC solution (100ng/μl 1:1 Trypsin (Promega) and LysC (FujiFilm) mix in 25mM NH4HCO3) and 21μl of 25mM NH4HCO3 (pH8.5) was added to 100µl final volume. Digestion was conducted overnight at 37°C. Reaction was terminated by acidification with 2.5% Trifluoroacetic acid (TFA) to 0.3% final.

#### NanoLC-MS/MS

Peptides were analyzed by nanoLC-MS/MS using the Agilent 1100 nanoflow system (Agilent) connected to hybrid linear ion trap-orbitrap mass spectrometer (LTQ-Orbitrap Elite™ (ThermoFisher)) equipped with an EASY-Spray™ electrospray source (held at constant 35°C). Chromatography of peptides prior to mass spectral analysis was accomplished using capillary emitter column (PepMap® C18, 3µM, 100Å, 150×0.075mm (ThermoFisher)) onto which 2µl of extracted peptides was automatically loaded. NanoHPLC system delivered solvents A: 0.1% (v/v) formic acid, and B: 99.9% (v/v) acetonitrile, 0.1% (v/v) formic acid at 0.50 µL/min to load the peptides (over a 30 minute period) and 0.3µl/min to elute peptides directly into the nano-electrospray with gradual gradient from 0% (v/v) B to 30% (v/v) B over 80 minutes followed by 5 minute fast gradient from 30% (v/v) B to 50% (v/v) B and concluded with a 5 minute flash-out from 50-95% (v/v) B. As peptides eluted from the HPLC-column/electrospray source survey MS scans were acquired in the Orbitrap with a resolution of 120,000 followed by CID-type MS/MS fragmentation of 30 most intense peptides detected in the MS1 scan from 350 to 1800 m/z; redundancy was limited by dynamic exclusion.

#### Data analysis

Elite acquired MS/MS data files were converted to mgf file format using MSConvert (ProteoWizard: Open-Source Software for Rapid Proteomics Tools Development). Resulting mgf files were used to search against Uniprot Rattus norvegicus proteome database (UP000002494, 10/06/2020 download, 31,681 total entries) along with a cRAP common lab contaminant database (116 total entries) using in-house Mascot search engine 2.7.0 (Matrix Science) with fixed Cysteine carbamidomethylation and variable Methionine oxidation plus Asparagine or Glutamine deamidation. Peptide mass tolerance was set at 10 ppm and fragment mass at 0.6 Da. Protein annotations, significance of identification and spectral based quantification was done with Scaffold software (version 4.11.0, Proteome Software Inc.). Peptide identifications were accepted if they could be established at greater than 88.0% probability to achieve an FDR less than 1.0% by the Scaffold Local FDR algorithm. Protein identifications were accepted if they could be established at greater than 6% probability to achieve an FDR less than 1.0% and contained at least 2 identified peptides. Protein probabilities were assigned by the Protein Prophet algorithm [92] that contained similar peptides and could not be differentiated based on MS/MS analysis alone were grouped to satisfy the principles of parsimony. Proteins sharing significant peptide evidence were grouped into cluster.

### Animal behavior

Rat plantar cutaneous mechanical sensitivity was assessed as previously reported [23]. Rats underwent 30uL intraplantar (footpad) injections of saline or treatment with undiluted Schwann cell collection media or p11 (0.9, 9, and 90ng) and were acclimated on top of a wire mesh for 1 hr, with the experimenter present for 30 mins of this period. Testing was performed at approximately the same time each day. Mechanical sensitivity threshold was determined with von Frey filaments (up-down method [93]) and values were analyzed after log transform [94]. Hypersensitivity to noxious force (hyperalgesia), which is selectively associated with aversion [95], was tested by needle prick [96]. The experimenter was blinded to treatment for all experiments. For baseline (BL) von Frey withdrawal threshold measurements, animals underwent testing within seven days of treatment administration.

### Statistical Analysis

Results were considered statistically significant when p < 0.05. All data were analyzed using GraphPad/Prism (Version 9.0.0). Statistical analyses used for each dataset are indicated within each figure legend. For dorsal root teased fiber single unit recording, data were analyzed using Chi-square for spontaneous activity per fiber bundle and an unpaired Student’s *t*-test for spontaneous activity per animal and firing rate. Mechanical sensitivity to graded von Frey stimulation was analyzed using a two-way repeated measures ANOVA, and sensitivity to needle analyzed using an unpaired Student’s *t*-test. For analysis of myelin pathology, data were analyzed using unpaired Student’s *t*-test. For current-clamp experiments, membrane and AP properties were analyzed using a one-way ANOVA for testing Schwann cell media effects on neuron function, and unpaired Student’s t-test for testing the effects of p11 on neuron function; current-evoked firing frequency was analyzed using a two-way repeated measures ANOVA. Percentage of spontaneously active cells was analyzed using Chi-square. For calcium imaging experiments, proportion of cells responding were analyzed using Chi-square, while response magnitude data were analyzed using a one-way ANOVA. For voltage-clamp experiments, current densities were measured using a two-way repeated measures ANOVA and maximum current density was analyzed using unpaired Student’s t-test. For nanoLC MS/MS data was analyzed using a Benjamini-Hochberg corrected two-tailed *t*-test. For ELISA experiments, data were analyzed using a two-way ANOVA. For von Frey and noxious needle tests, data were analyzed using a two-way repeated measures ANOVA. Bonferroni post hoc corrections were used for significant ANOVAs. Fisher’s exact tests were used for significant Chi-square tests.

### Study Approval

All protocols were in accordance with National Institutes of Health guidelines and were approved by the Institutional Animal Care and Use Committee at the Medical College of Wisconsin.

## Acknowledgments

We would like to thank Clive Wells for help with processing light microscopy and TEM samples. We show appreciation for Greg Sabat and the University of Wisconsin Mass Spectrometry Core Facility for conducting mass spectrometry experiments. We would like to thank Dr. Stephanie Shiffka for advice on interpretation of mass spectrometry data. We would like to thank James J. Miller and Carly A. Mascari for harvesting DRG tissue, as well as Chris Duris and Suresh Kumar at the Children’s Research Institute’s Imaging Core at the Medical College of Wisconsin for processing and imaging H&E slides. We would like to thank Katelyn Sadler for advice and expertise on preparing the manuscript. All figure graphics and the graphical abstract were created using BioRender.com.

## Author Contributions

TBW, CLS conceptualized the experiments. TBW, DC, VLE, OI, BP developed the meth-odology. TBW, DC, EKP performed the experiments. TBW and DC analyzed the data. TBW created the original draft manuscript. TBW and BSD created figure visualizations. All authors contributed to the editing of the paper. Supervision was provided by CLS, BP, QH, NMD. Funding for this work was acquired by CLS, NMD, TBW.

