## Supplemental Figures for "Schwann cell release of p11 induces sensory neuron hyperactivity in Fabry disease"

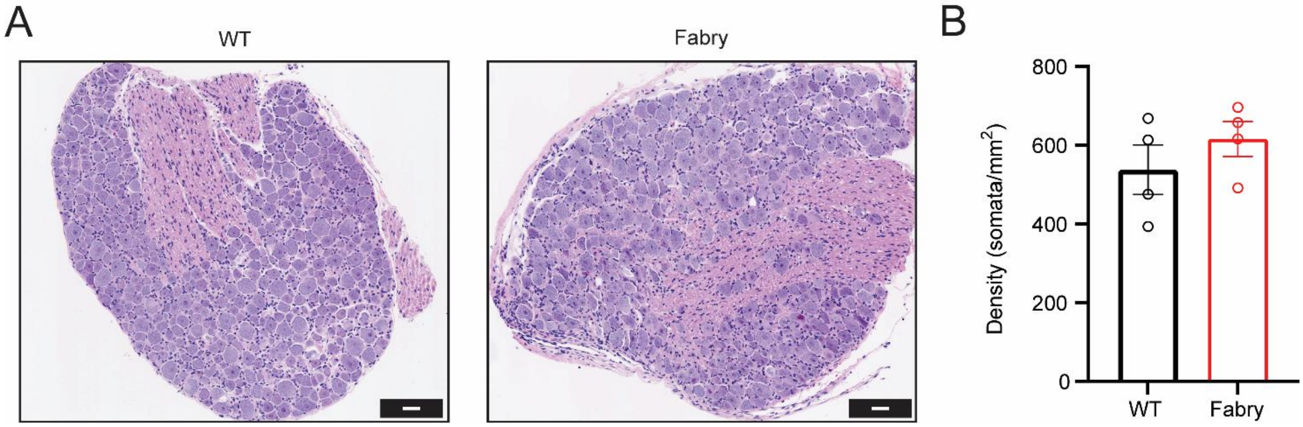


Figure 1S: Lumbar DRG isolated from Fabry and WT rats exhibit similar neuron soma density. A) Representative cross-sections of lumber (L1) DRG from WT and Fabry rats, stained with hematoxylin and eosin (H&E), scale bar 50µm. B) Both genotypes exhibit a similar density of somata per DRG, Values reported as mean ± SEM. (B) unpaired *t*-test. n = 4 animals per genotype, one DRG cross section analyzed per animal.

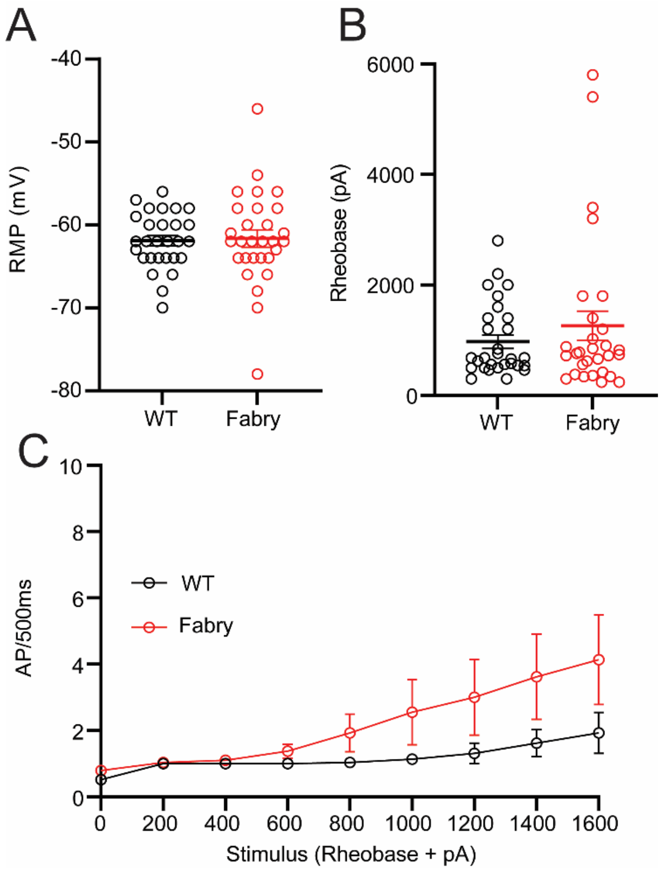


Figure 2S: Large diameter DRG neuron somata isolated from Fabry and WT rats exhibit similar current-evoked excitability. A) Both Fabry and WT large diameter (>32 µm) exhibit similar resting membrane potentials and B) rheobase. C) Both genotypes exhibit similar firing frequency to suprathreshold current stimulation starting from rheobase to rheobase + 1,600 pA. n = 29 neurons from 8 animals per genotype. Values reported as mean ± SEM. (A, B) unpaired *t*-test, (C) two-way repeated measures ANOVA. RMP = resting membrane potential, AP = action potential.

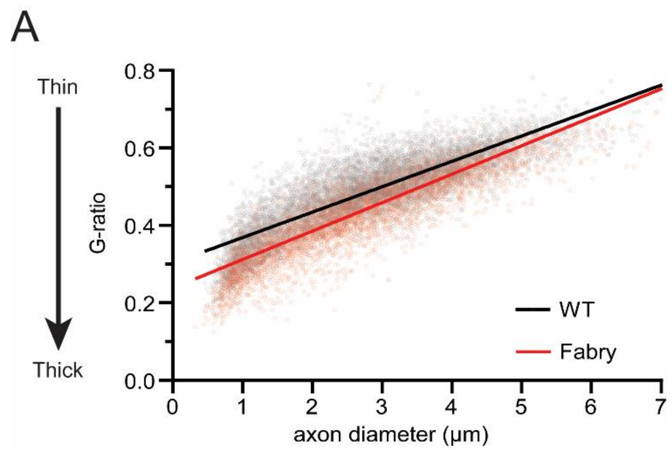


<u>Figure 3S: Fabry and WT axon G-ratio derived from tibial nerve fascicles plotted against axon diameter.</u> Simple linear regression of axon g-ratios plotted against axon diameter from cross-sectioned tibial nerves of WT and Fabry rats. n = g-ratio from 7,000 – 8,000 axons derived from cross-sectioned tibial nerves of 6 animals per genotype.

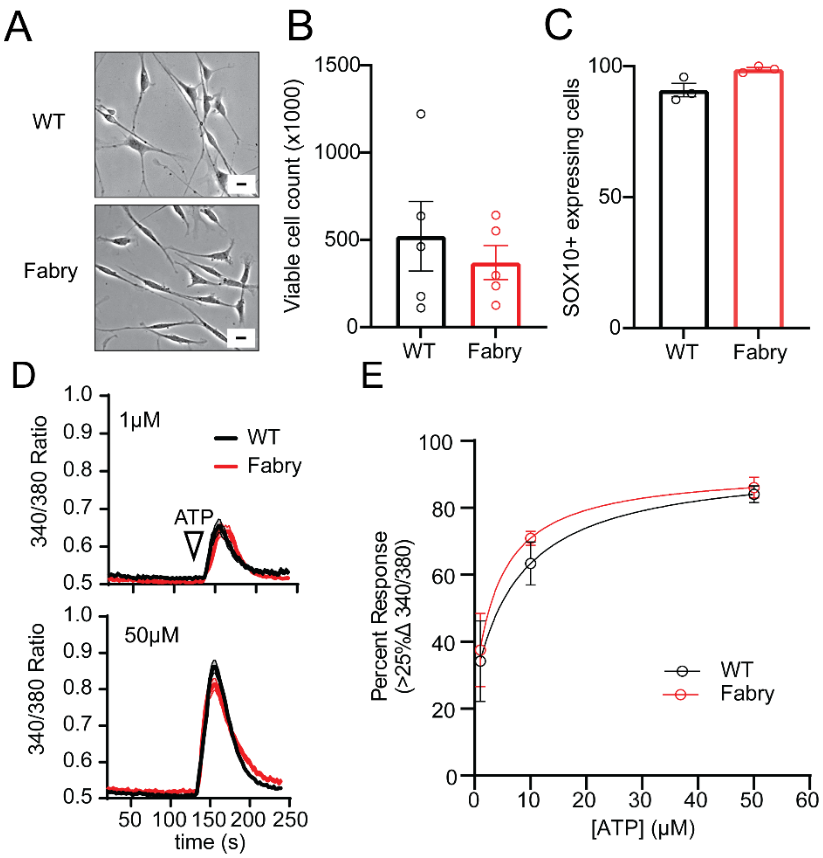


<u>Figure 4S: Fabry and WT Schwann cells exhibit similar cell health.</u> A) Representative light microscopy images of Fabry and WT Schwann cells in culture, scale bar 10 μm. B) WT and Fabry Schwann cell cultures possessed similar cell viability via Trypan blue analysis two days after plating. C) Cultured Schwann cells from both genotypes exhibited high purity two days after plating as quantified by immunofluorescent imaging with the Schwann cell marker SOX10 (each dot represents the averaged SOX10+ cell percentage gathered from 3 animals per genotype). D) Aggregated traces depicting the mean ± SEM of WT and Fabry Schwann cells exposed to 1 or 50 μM ATP for 10s. E) Calcium imaging analysis revealed a similar concentration-dependent response to ATP in both genotypes. (B, C) each dot represents per animal measurement. (D, E) n = 200 Schwann cells per group from 3 animals. Values reported as mean ± SEM.

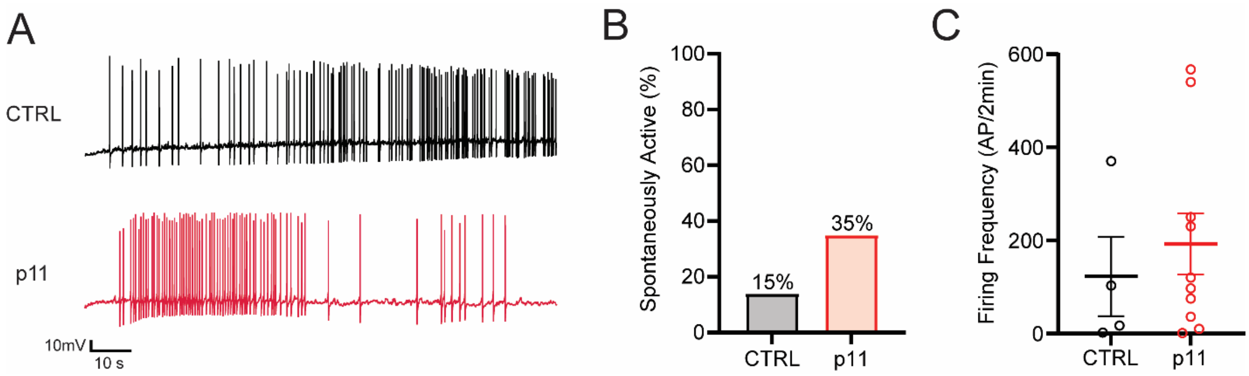


<u>Figure 5S: Naïve neurons exposed to overnight incubation of p11 exhibit a non-significant trend towards increased spontaneous activity.</u> A) Representative spontaneous traces of naïve DRG neurons exposed to 100 ng/mL of p11 or CTRL. B) A trending, but not significant, number of neurons exposed to p11 exhibited spontaneous activity compared to CTRL neurons. n = 28 neurons per treatment from 7 animals C) Spontaneous firing frequency of DRG neurons were similar between treatments. (B) reported as mean, (C) reported as mean ± SEM. (B) $\chi^2$, p = 0.12, (C) unpaired *t*-test.

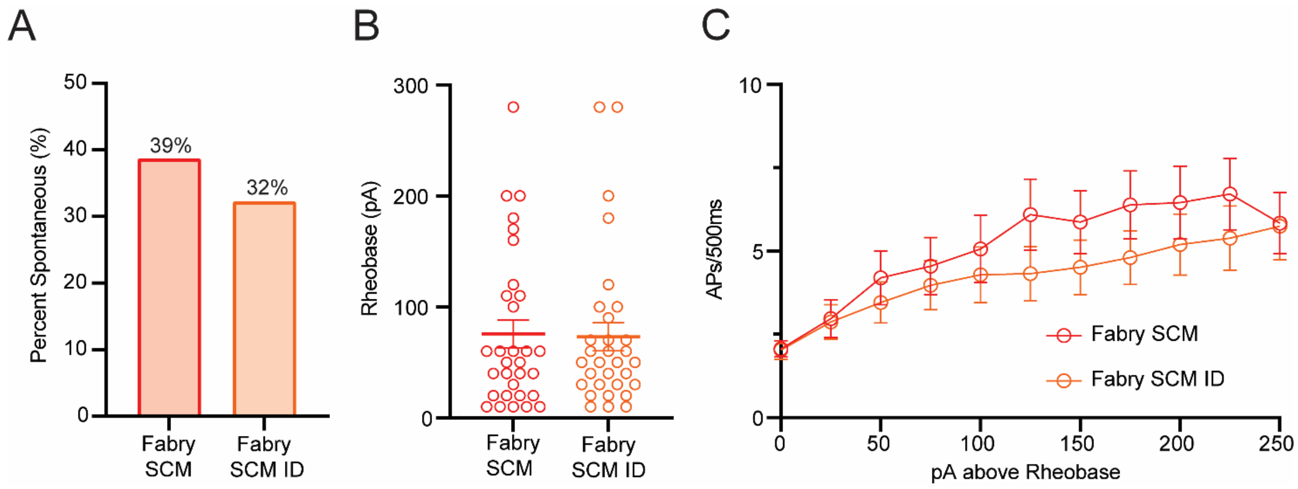


Figure 6S: Naïve neurons treated with either Fabry SCM or Fabry SCM ID exhibit similar spontaneous activity and current-evoked excitability. A) A similar proportion of DRG neuron somata exhibit spontaneous activity when exposed to Fabry SCM or Fabry SCM with immunodepleted p11 (ID) overnight. B) DRG neurons exposed to either treatment exhibit a similar rheobase and C) current-evoked firing frequency. n = 31 neurons per treatment from 8 animals. (A) reported as mean and (B) mean ± SEM. (A) $\chi^2$, (B) unpaired *t*-test, (C) two-way repeated measures ANOVA. Fabry SCM = Fabry Schwann cell conditioned media. Fabry SCM ID = Fabry Schwann cell conditioned media with immunodepleted p11.

| Small Diameter (≤ 32um) | | |
|---|---|---|
| | **WT** | **Fabry** |
| Cell Number | 72 | 70 |
| RMP (mV) | -57.7 ± 0.9 | -56.9 ± 1.1 |
| **Spontaneous Activity** | | |
| Cell Number | 34 | 33 |
| Cell Size (μm) | 24.02 ± 0.4 | 24.30 ± 0.6 |
| Capacitance (pF) | 43.75 ± 2.9 | 37.6 ± 2.5 |
| Percent Spontaneous (%) [b] | 11.7% | 33.3% * |
| Firing Frequency (AP/2min) | 424.3 ± 289.9 | 201.5 ± 80.92 |
| **Current-Evoked Excitability** | | |
| Cell Number | 38 | 37 |
| Cell Size (μm) | 25.3 ± 0.5 | 25.07 ± 0.6 |
| Capacitance (pF) | 37.7 ± 1.9 | 37.9 ± 2.3 |
| Input Resistance (mΩ) | 561.3 ± 48.9 | 487.2 ± 57.6 |
| Rheobase (pA) [a] | 300.0 ± 39.1 | 202.0 ± 22.8 * |
| AP Threshold (mV) | -18.7 ± 1.9 | -21.0 ± 1.7 |
| AP Amplitude (mV) | 42.8 ± 2.1 | 40.8 ± 2.1 |
| AP Half-width (ms) | 1.6 ± 0.1 | 1.5 ± 0.1 |

**Table 1: Passive and current-evoked membrane properties of small diameter DRG neurons isolated from WT or Fabry rats.** Unpaired *t*-test = [a], * $p < 0.05$. $\chi^2$ main effect of treatment = [b], Fisher's exact post hoc comparisons, * $p < 0.05$. RMP: resting membrane potential, AP: action potential. Reported as Mean ± SEM.

| Large Diameter (> 32um) | | |
|---|---|---|
| **Current-Evoked Excitability** | | |
| | **WT** | **Fabry** |
| Cell Number | 29 | 29 |
| Cell Size (μm) [a] | 39.1 ± 1.0 | 43.9 ± 1.7 * |
| Capacitance (pF) | 92.12 ± 9.0 | 103.9 ± 10.5 |
| RMP (mV) | -61.9 ± 0.6 | -61.6 ± 1.1 |
| Input Resistance (mΩ) | 127.2 ± 18.3 | 185.0 ± 30.8 |
| Rheobase (pA) | 976.6 ± 121.6 | 1264.0 ± 264.9 |
| AP Threshold (mV) | -29.1 ± 2.1 | -25.8 ± 2.7 |
| AP Amplitude (mV) | 47.0 ± 2.1 | 47.1 ± 1.8 |
| AP Half-width (ms) | 0.95 ± 0.09 | 1.17 ± 0.19 |

**Table 2: Passive and current-evoked membrane properties of large diameter DRG neurons isolated from WT or Fabry rats.** Unpaired *t*-test = [a], * $p < 0.05$. RMP: resting membrane potential, AP: action potential. Reported as Mean ± SEM.

| | CTRL SCM | WT SCM | Fabry SCM |
|---|---|---|---|
| Cell Number | 51 | 59 | 60 |
| RMP (mV) [a] | -56.1 ± 1.1 | -54.8 ± 1.1 | -51.0 ± 1.1 ** # |
| **Spontaneous Activity** | | | |
| Cell Number | 23 | 28 | 29 |
| Cell Size (μm) | 25.5 ± 0.3 | 24.2 ± 0.4 | 24.30 ± 0.8 |
| Capacitance (pF) | 34.8 ± 2.1 | 33.4 ± 2.8 | 28.3 ± 2.5 |
| Percent Spontaneous (%) [b] | 4% | 18% | 52% ** # |
| Firing Frequency (AP/2min) | 5 ± 0 | 484 ± 149.3 | 295.8 ± 137.2 |
| **Current-Evoked Excitability** | | | |
| Cell Number | 28 | 31 | 31 |
| Cell Size (μm) | 25.4 ± 0.3 | 24.6 ± 0.6 | 24.2 ± 0.5 |
| Capacitance (pF) | 34.0 ± 2.8 | 33.2 ± 2.8 | 25.9 ± 2.1 |
| Input Resistance (mΩ) | 411.4 ± 39.5 | 501.3 ± 49.4 | 615.9 ± 82.1 |
| Rheobase (pA) | 291.4 ± 87.7 | 206.1 ± 23.8 | 133.2 ± 18.0 |
| AP Threshold (mV) | -19.6 ± 1.9 | -20.3 ± 1.7 | -22.6 ± 1.3 |
| AP Amplitude (mV) | 43.9 ± 1.3 | 42.1 ± 2.1 | 41.8 ± 1.9 |
| AP Half-width (ms) | 1.62 ± 0.1 | 1.70 ± 0.09 | 1.67 ± 0.09 |

**Table 3. Membrane properties of DRG neurons isolated from naïve rats following overnight treatment of WT, Fabry, or unconditioned (CTRL) Schwann cell media (SCM)**. One-way ANOVA main effect of treatment = [a] ; Bonferroni post-hoc tests. $\chi^2$ main effect of treatment = [b] ; Fisher's exact post hoc comparisons. CTRL vs. Fabry = *, ** $p < 0.01$, CTRL vs. Fabry #, # $p < 0.05$). RMP: resting membrane potential, AP: action potential. Reported as Mean ± SEM.

| Upregulated Protein Release (Fabry) | | | |
|---|---|---|---|
| **Protein** | **WT (Normalized Spectra)** | **Fabry (Normalized Spectra)** | **Fold Change (Fabry)** |
| p11 (S100A10) | 0.0 ± 0.0 | 0.93 ± 0.27 ** | ∞ |
| GSDMA | 0.0 ± 0.0 | 0.94 ± 0.31 * | ∞ |
| TGM1 | 0.14 ± 0.14 | 1.69 ± 0.57 * | 11 |
| TPI1 | 0.63 ± 0.27 | 1.53 ± 0.25 * | 2.4 |
| **Downregulated Protein Release (Fabry)** | | | |
| **Protein** | **WT (Normalized Spectra)** | **Fabry (Normalized Spectra)** | **Fold Change (Fabry)** |
| ENO2 | 3.59 ± 0.16 | 1.69 ± 0.33 *** | 0.5 |
| PKM | 14.13 ± 1.29 | 7.21 ± 1.50 *** | 0.5 |
| GPI | 7.18 ± 1.04 | 3.93 ± 0.58 * | 0.5 |
| TAGLN2 | 2.12 ± 0.46 | 0.76 ± 0.20 * | 0.5 |
| EEF2 | 10.38 ± 1.54 | 5.51 ± 1.11 * | 0.4 |
| PSMA6 | 1.16 ± 0.35 | 0.21 ± 0.21 * | 0.5 |
| EIF5A | 0.85 ± 0.36 | 0.0 ± 0.0 * | 0 |

**Table 4. Significantly different protein concentrations between WT and Fabry Schwann cell media as identified by NanoLC-MS/MS analysis.** Benjamini-Hochberg corrected two-tailed *t*-test, * $p < 0.05$, ** $p < 0.01$, ***$p < 0.001$. Reported as Mean ± SEM.

| | CTRL | p11 |
|---|---|---|
| Cell Number (total) | 28 | 28 |
| Cell Size (μm) | 26.7 ± 0.4 | 25.3 ± 0.4 |
| Capacitance (pF) | 35.4 ± 2.1 | 31.7 ± 1.6 |
| RMP (mV) | -58.6 ± 1.4 | -52.1 ± 1.6 ** |
| **Spontaneous Activity** | | |
| Percent Spontaneous (%) | 14% | 35% |
| Firing Frequency (AP/2min) | 123.0 ± 85.3 | 207.6 ± 65.7 |
| **Current-Evoked Excitability** | | |
| Input Resistance (mΩ) | 466.3 ± 38.8 | 535.7 ± 40.1 |
| Rheobase (pA) | 174.6 ± 25.8 | 92.1 ± 22.8 * |
| AP Threshold (mV) | -15.4 ± 1.7 | -15.1 ± 1.9 |
| AP Amplitude (mV) | 43.5 ± 1.2 | 41.2 ± 1.3 |
| AP Half-width (ms) | 1.94 ± 0.15 | 1.90 ± 0.11 |

**Table 5. Passive and current-evoked membrane properties of DRG neurons isolated from naïve rats following overnight treatment of 100ng/mL p11 or vehicle (CTRL).** Unpaired *t*-test, * $p < 0.05$, ** $p < 0.01$. RMP: resting membrane potential, AP: action potential. Reported as Mean ± SEM.

| | Fabry SCM | Fabry SCM ID |
|---|---|---|
| Cell Number (total) | 31 | 31 |
| Cell Size (µm) | 23.2 ± 0.3 | 23.5 ± 0.4 |
| Capacitance (pF) | 24.02 ± 1.0 | 25.8 ± 1.2 |
| RMP (mV) | -49.1 ± 1.3 | -54.6 ± 1.5 ** |
| **Spontaneous Activity** | | |
| Percent Spontaneous (%) | 38.7% | 32.3% |
| Firing Frequency (AP/2min) | 222.8 ± 97.3 | 49.4 ± 33.2 |
| **Current-Evoked Excitability** | | |
| Input Resistance (mΩ) | 817.6 ± 91.0 | 620.1 ± 41.8 |
| Rheobase (pA) | 75.81 ± 12.6 | 73.2 ± 12.7 |
| AP Threshold (mV) | -16.8 ± 1.9 | -17.8 ± 1.8 |
| AP Amplitude (mV) | 39.8 ± 2.1 | 43.0 ± 2.1 |
| AP Half-width (ms) | 1.9 ± 0.09 | 1.8 ± 0.09 |

**Table 6: Passive and current-evoked membrane properties of DRG neurons isolated from naïve rats following overnight treatment of Fabry Schwann cell media (SCM) or Fabry SCM with immunodepleted (ID) p11.** Unpaired *t*-test, ** $p < 0.01$. RMP: resting membrane potential, AP: action potential. Reported as Mean ± SEM.
